# The diel structure and robustness of plant-pollinator networks are driven by the shifting influences of plant traits

**DOI:** 10.64898/2026.08.05.742987

**Authors:** Sebastián Montoya-Bustamante, Casper J. van der Kooi, Vincent Grognuz, Colin Fontaine, Eva Knop

## Abstract

1. Species interactions are increasingly recognised as temporally dynamic. For plant-pollinator networks, evidence shows that interactions vary not only seasonally but also over the diel cycle. However, we still know little about what determines their diel structure, whether this structure is important for robustness to species loss, and which plant traits are associated with the roles plants play in this structure. These gaps are fundamental, because knowing what shapes networks over the diel cycle is required to predict the effects of global change drivers, such as light pollution, that may shift the timing of interactions.
2. Using 22 plant-pollinator networks sampled over morning, afternoon, and night, we addressed these gaps by applying a multilayer framework to characterise their diel structure, link it to robustness, and test which plant traits are associated with plant roles across diel periods (participation, versatility) and within them (centrality).
3. Diel structure was non-random: interactions were segregated among diel periods yet integrated through plants visited across the diel cycle. Networks were more robust to simulated species loss when interactions were on average more evenly distributed across periods and when plants were more strongly interconnected among periods, although this benefit diminished when both properties were high simultaneously. The association between plant traits and their roles shifted with the temporal scale. Across diel periods, structural traits were the stronger predictors: taller plants were visited more evenly across the diel cycle (higher participation), whereas shorter plants shared pollinators with plants from multiple periods (higher versatility), potentially mediating indirect effects among them. Within diel periods, floral visual cues became more influential, with achromatic contrast the most consistent predictor: at night, plants with brighter flowers were well visited within the period, sharing pollinators with other plants of that period (higher centrality).
4. These findings identify diel structure as a functional axis of network organisation and indicate that plant-pollinator networks are assembled hierarchically: structural traits set a baseline across the diel cycle, whereas light conditions determine which traits matter within periods, ultimately defining distinct temporal pathways vulnerable to global change.

## 1 Introduction

Plants and their pollinators are entangled in complex interaction networks, which are shaped by species’ traits, behaviours, and environmental conditions that vary across space and time (Junker *et al*. 2013; Olesen *et al*. 2007; Schwarz *et al*. 2020). As a result, plant-pollinator networks are inherently dynamic (Schwarz *et al*. 2020). Temporal dynamics are increasingly recognised as essential for understanding network structure and function, yet remain critically understudied (CaraDonna *et al*. 2021; Schwarz *et al*. 2020). Most research has focused on seasonal dynamics (reviewed by CaraDonna et al. 2021), and only recently has attention shifted towards diel dynamics (Knop *et al*. 2017; Schwarz *et al*. 2021; Souza *et al*. 2022; Teng *et al*. 2024). Diel patterns in plant-pollinator networks are likely driven by multiple factors, including pollinator activity and plant traits (Bloch *et al*. 2017; Borges *et al*. 2016; Knop *et al*. 2018). Understanding these patterns is becoming urgent, because global change drivers such as light pollution may shift the timing of interactions (Briolat *et al*. 2021; Grognuz *et al*. 2026), and predicting their consequences requires knowing how interactions are organised over the diel cycle in the first place. However, we still lack an understanding of (1) whether plant-pollinator networks indeed show a non-random *diel structure* (i.e., whether interactions are temporally compartmentalised over the diel cycle), (2) whether diel structure is important for network *robustness* (i.e., the capacity of a network to sustain function despite species loss), and (3) whether diel structure is associated with plant traits.

The diel structure of plant-pollinator networks is likely compartmentalised over the diel cycle. This expectation stems from pronounced diel variation in pollinator activity, driven by circadian clocks, abiotic conditions, and light (Bloch *et al*. 2017; Borges *et al*. 2016). Different pollinator groups exhibit distinct activity peaks (Knop *et al*. 2018; Xu *et al*. 2021; Zoller *et al*. 2020). For example, activity by dipterans tends to peak in the morning, hymenopterans in the afternoon, and lepidopterans at night (Herrera 1990; Knop *et al*. 2018). Even within daytime, some taxa show bimodal activity with distinct morning and afternoon peaks (Xu *et al*. 2021; Zoller *et al*. 2020). As a result, pollinator community composition may differ among diel periods (Gårdman *et al*. 2026). Morning, afternoon, and night therefore provide a biologically grounded division of the diel cycle. Compartmentalisation among these periods could influence network robustness, because perturbations spread more readily in less compartmentalised structures (Gilarranz *et al*. 2017; Grilli *et al*. 2016; May 1972; Stouffer & Bascompte 2011). However, the relationship between network robustness and the diel structure of plant-pollinator networks remains unresolved.

Besides pollinator activity, plant traits may also be associated with a network’s diel structure, yet the influence of plant traits across temporal scales is unclear. We hypothesise that plant traits operate through distinct pathways at different temporal scales. *Across diel periods*, structural traits that remain largely constant and do not depend on light conditions, such as floral (morphological) complexity and plant height, may set a baseline for potential interactions. As an example, taller plants and less complex flowers are better accessible to a broad range of visitors, whereas smaller plants and complex flowers tend to be accessible to specific visitors (Engel & Irwin 2003; Junker 2012; Junker *et al*. 2013; Krishna & Keasar 2018). *Within diel periods*, by contrast, light-associated traits may become more important. These include traits that enhance pollination success, such as floral opening, scent emission, and reward availability, which often follow plants’ circadian clocks (Bloch *et al*. 2017; van Doorn & Kamdee 2014; van Doorn & van Meeteren 2003; Kendall & Nicholson 2025; Roemer *et al*. 2022), and visual cues whose relevance depends on the ambient light (Kelber *et al*. 2003; van der Kooi *et al*. 2019). Among these visual cues, flower colour and achromatic (brightness) contrast both determine flower visibility to diurnal and nocturnal flower visitors (Kelber *et al*. 2003; van der Kooi *et al*. 2019). Under low light conditions, achromatic contrast is expected to be particularly important for nocturnal visitors (van der Kooi & Kelber 2022).

Testing whether plant traits are associated with the diel structure of plant-pollinator networks requires analytical tools that capture temporal organisation. Multilayer network analysis offers a powerful approach (Battiston *et al*. 2014; De Domenico *et al*. 2015; Pilosof *et al*. 2017): interactions occurring in morning, afternoon, and night can be represented as distinct layers, allowing us to quantify plants’ roles *across* and *within* layers (De Domenico 2022; Hutchinson *et al*. 2019; Pilosof *et al*. 2017). This dual perspective maps directly onto our hypothesis: On the one hand, *across* layers, we can analyse plants’ roles through their *participation*, which quantifies how evenly a plant’s interactions are distributed across diel periods (Battiston *et al*. 2014; Guimerà & Nunes Amaral 2005), and their *versatility*, which quantifies a plant’s importance for network connectivity across multiple periods (De Domenico 2022; De Domenico *et al*. 2015). Averaged over all plants in a network, participation and versatility respectively map onto two aspects of temporal compartmentalisation: segregation vs. evenness in the distribution of interactions, and fragmentation vs. integration of the diel periods. On the other hand, *within* layers, plants’ roles can be assessed by estimating their *centrality*, which quantifies a plant’s importance for network connectivity within specific periods (Maia *et al*. 2019; Martín González *et al*. 2010). Together, *participation*, *versatility*, and *centrality* allow us to characterise diel structure and test whether plant traits are associated with the roles plants occupy within it.

In this work, we aimed at understanding the drivers and consequences of the diel structure of plant-pollinator networks. We asked: (1) Given a partition into morning, afternoon, and night, is the diel structure of plant-pollinator networks more temporally compartmentalised than expected by chance? (2) What is the relationship between diel structure and network robustness? (3) How are plant structural traits and floral visual cues associated with the roles plants occupy *across* and *within* diel periods? Our findings allowed us to reveal hidden drivers of the structure and robustness of ecological networks.

## 2 Material and methods

### 2.1 Experimental design

Between 2014 and 2017, during spring and summer, we selected 15 independent ruderal meadows across the Prealps of Switzerland (Appendix S1, Table S1). All sites were comparable in vegetation, with *Cirsium oleraceum* (Asteraceae) usually being the most abundant. Not all sites were visited in all years and the number of sites visited per year was variable (Table S1). Further details of the sites are available in Knop *et al*. (2017, 2018), and Giavi *et al*. (2020, 2021).

### 2.2 Sampling plant-pollinator interactions

Plant-pollinator interactions were sampled using sweep nets at three different diel periods, which were determined based on the relative position of the sun in the sky: (1) “morning”, from sunrise to solar noon; (2) “afternoon”, from solar noon to sunset, and (3) “night”, from sunset to sunrise (Appendix S1). To objectively determine the limits of these periods, we used the package SUNCALC v0.5.1 of R (Thieurmel & Elmarhraoui 2025). During 2015 and 2017, samples were taken only at night, whereas during 2014 and 2016, samples were taken on at least two periods, always including the night (Table S2).

The interactions of a plant species *j* with a given pollinator species *i* were quantified as the number of visits of *i* in which they were in contact with the reproductive organs of the flowers of *j*; a standard approach in plant-pollinator network studies (e.g., CaraDonna & Waser (2020), Schwarz et al.(2021)). However, not all visitors were considered as pollinators. We classified pollinators depending on the morphology of their mouthparts, which is a good predictor of their diet (Krenn 2019; Krenn *et al*. 2005). Specifically, only those visitors whose mouthparts are adapted to access typical floral rewards such as nectar and pollen were included. This criterion allowed us to exclude groups like dermapterans, orthopterans, and some coleopterans which often feed on and damage floral reproductive organs (Karolyi 2019; Krenn *et al*. 2005) (Table S3).

### 2.3 Flower abundance

A flower assessment was performed during each sampling day, where occurring plant species were identified. For each plant species *j*, flower abundance was estimated as the sum of the floral units of *j* during the whole sampling. A floral unit was defined as the area of 5 cm^2^ occupied by open flowers of a given species. On a few occasions, the quadrants for the flower assessment did not exactly match the interaction sampling quadrants, leaving us with some plant species without a flower abundance estimation. For these cases, we assigned to those plants the same flower abundance of the least abundant plant recorded in the same site and year.

### 2.4 Compiling information on plant traits

To address our questions, we compiled information on two different types of plant traits (Table S4): (1) structural traits and (2) visual cues.

#### 2.4.1 Structural traits

We focused on two structural traits: (1) plant (maximum) height, as a proxy for flower height above/in vegetation, and (2) floral (morphological) complexity.

##### 2.4.1.1 Plant height

Data on plant height was obtained from Lauber et al. (Lauber *et al*. 2018) via InfoFlora (https://www.infoflora.ch/), and complemented with data from FloraVeg.EU (Chytrý *et al*. 2024). Values of plant height exceeding 2 m were kept at that value, because no flower was sampled over that height. We were able to obtain data on plant height for 117 species.

Although plant height could be considered as a visual cue (Michelot-Antalik *et al*. 2025), we kept it in this category because we do not expect its effect to change with light conditions.

##### 2.4.1.2 Floral complexity

Floral complexity is a difficult trait to measure because it encompasses multiple floral traits associated with the accessibility of pollinators to floral rewards, that is, there are many ways to be a complex flower (Keasar 2020). Therefore, for each flower, we calculated the floral complexity index (FCI), defined as the sum of the “complexity weights” of its floral traits (Stefanaki *et al*. 2015). The traits used to calculate the FCI were: (1) floral shape, (2) floral depth, (3) floral symmetry, (4) corolla segmentation, and (5) functional reproductive unit. Higher values of FCI indicate lower accessibility to floral resources. Trait weights and their assignment are detailed in Appendix S2, and FCI values for each plant in Table S5.

#### 2.4.2 Visual cues

We focused on two floral visual cues: colour contrast and achromatic contrast. To calculate these two types of visual contrast, we obtained data on reflectance spectra from the Floral Reflectance Database (FReD, Arnold *et al*. (2010)) and from published work (van der Kooi *et al*. 2016b, a) (Table S6). Flower colours were interpreted with an “insect-subjective view”, by deploying the hexagon vision model (Chittka 1992), using honeybee spectral sensitivity, a D65 illuminant and a green (leaf) background. Colour contrast was calculated as the Euclidian distance between a flower stimulus and the centre of the colour space (Chittka 1992; Kelber *et al*. 2003; van der Kooi & Spaethe 2025). Achromatic contrast was calculated as the absolute difference in the long-wavelength photoreceptor excitation between the flower stimulus and background (Giurfa *et al*. 1996; Spaethe *et al*. 2001). Out of 119 plant species included in our analysis, we were able to obtain achromatic and colour contrast values for 79.

### 2.5 Network analysis

We built 22 plant-pollinator networks, one per sampled site and year (Table S2). Fourteen networks spanned multiple diel periods; the rest only night. For each network, interaction data from each diel period were initially organized into bi-adjacency matrices (hereafter “period matrices”). Pollinator species were placed in rows and plant species in columns, with cell values representing the interaction frequency—the number of visits 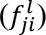between pollinator *i* and plant *j* in period *l*. All analyses were run in R (R Core Team 2023). All statistical models were made using the packages GLMMTMB v1.1.13 (Brooks *et al*. 2017), DHARMA v0.4.7 (Hartig 2022), and PERFORMANCE v0.15.02 (Lüdecke *et al*. 2021).

#### 2.5.1 Given a partition into morning, afternoon, and night, is the diel structure of plant-pollinator networks more temporally compartmentalised than expected by chance?

To assess the temporal compartmentalisation in the diel structure of plant-pollinator networks, we analysed the 14 networks sampled across multiple diel periods and characterised diel structure from the plant perspective using two complementary metrics (Fig. 1): *mean participation* (*P̅*) and *mean versatility* (*V̅*). These metrics capture distinct aspects of temporal compartmentalisation as explained below.

**Figure 1.**
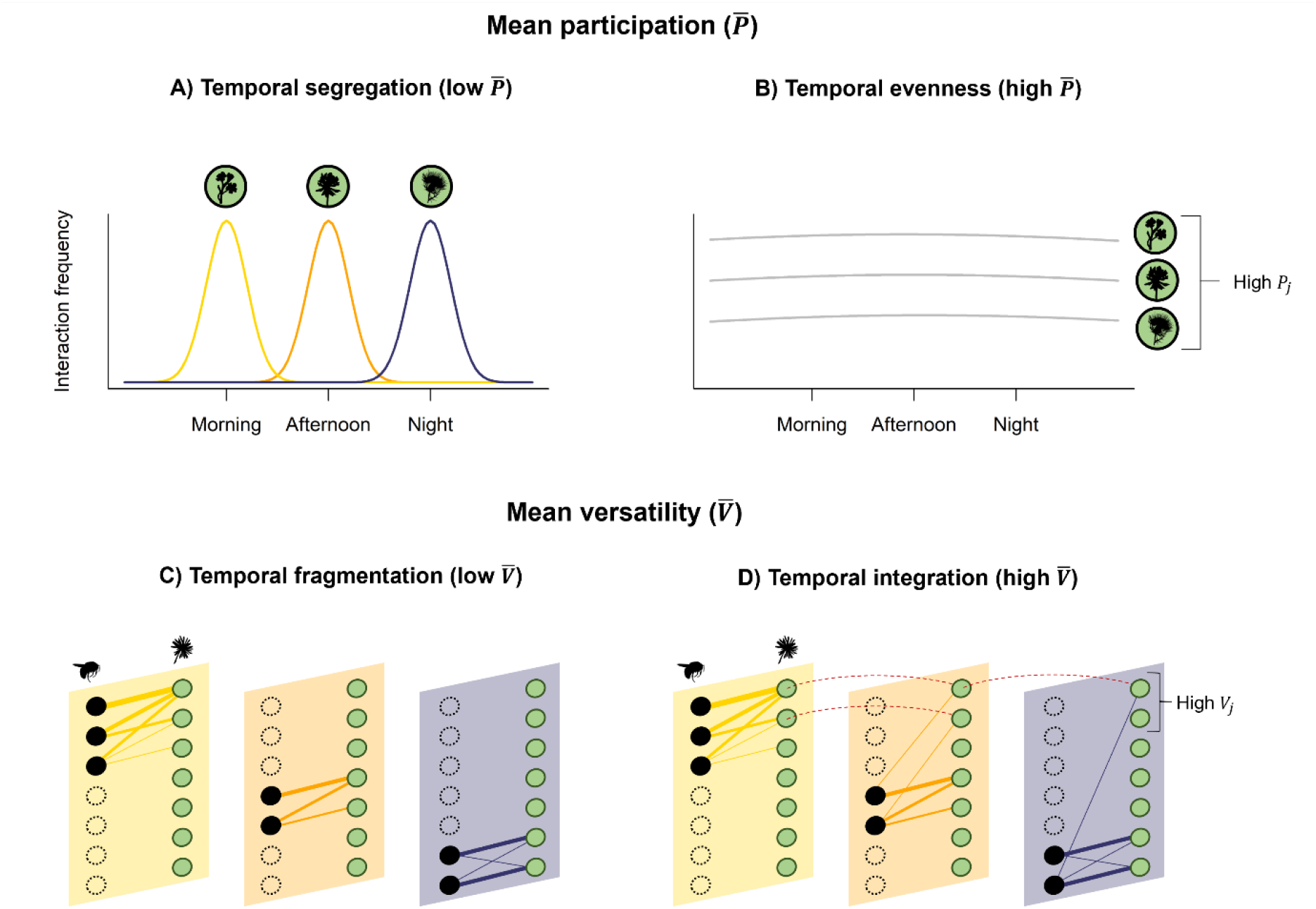
Mean participation and mean versatility offer related yet complementary insights about temporal compartmentalisation in the diel structure of networks. Participation (*P_j_*) quantifies how evenly the interactions of a plant *j* are distributed across diel periods. Mean participation (*P̅*) is low when plant interactions are concentrated within a single diel period, indicating temporal segregation (A). Conversely, mean participation is high when interactions are evenly distributed across all periods, indicating temporal evenness (B). Versatility (*V_j_*) quantifies how important a plant *j* is for maintaining cross-period connectivity. Mean versatility (*V̅*) is low when most plants are interconnected within isolated diel periods, indicating temporal fragmentation (C). Conversely, mean versatility is high when plants bridge different diel periods, indicating temporal integration (D). Note that a plant can exhibit low *P_j_* and high *V_j_* if it bridges multiple diel periods but most of their interactions are concentrated within a single period, as it is shown for the top plants in panel D, whose interactions are concentrated in the morning. Black and green circles represent pollinators and plants, respectively. Solid lines between them represent interactions and frequency is indicated by their width. Dashed lines indicate that a plant is bridging the diel periods. This is an illustration for explanatory purposes only.

##### 2.5.1.1 Mean participation

Mean participation quantifies *temporal evenness* by capturing how uniformly plants’ interactions are distributed across diel periods (Fig. 1A—B). Specifically, *P̅* is the average of plant-level participation scores (*P_j_*; the cross-period interaction evenness of plant *j*) in a network. (Battiston *et al*. 2014; Guimerà & Nunes Amaral 2005). Participation is calculated as 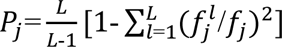, where 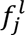 is the interaction frequency of plant *j* in diel period *l* and *f_j_* is the total interaction frequency of *j* across all *L* diel periods (Battiston *et al*. 2014). *P_j_* ranges from 0.0 (all interactions of plant *j* are segregated in one diel period) to 1.0 (interactions of plant *j* are evenly distributed across diel periods). Likewise, low values of mean participation over all plants in a network (*P̅*) indicate *temporal segregation*, where plants consistently interact in specific few diel periods (Fig. 1A). Conversely, high values indicate *temporal evenness*, where plant interactions are distributed uniformly across periods (Fig. 1B).

##### 2.5.1.2 Mean versatility

Mean versatility quantifies *temporal integration* by estimating how well interconnected diel periods are in a network (Fig. 1C—D). Specifically, *V̅* is the average of plant-level versatility scores (*V_j_*; the importance of plant *j* for cross-period connectivity by sharing species with plants from multiple periods) in a network (De Domenico *et al*. 2015). Details on the calculation of versatility are found in Appendix S3. The values of versatility in each network were normalized to range from 0.0 (low cross-period importance) to 1.0 (high cross-period importance). Therefore, low values of mean versatility over all plants in a network (*V̅*) indicate *temporal fragmentation* (Fig. 1C), where the diel periods operate as disconnected temporal clusters. Conversely, high values indicate *temporal integration*, where plants consistently bridge multiple diel periods (Fig. 1D).

##### 2.5.1.3 Null models

To evaluate whether the observed temporal patterns in our networks differed from random expectations, we constructed three temporal null models (Fig. S1). Null model 1 preserved overall plant interaction frequency, randomly redistributing interactions among all pollinators and periods. Null model 2 added a constraint by also preserving period-specific interaction frequency. Null model 3 further constrained the system by maintaining period-specific pollinator interaction frequency. For each of the 20 networks, we generated 1,000 null networks per model, for a total of 60,000. Then, we calculated their mean participation and mean versatility, and estimated the P value for each observed network as 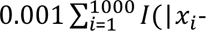 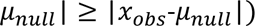, where *x_i_* is *i*-th value from the null distribution, *μ_null_* is the mean of the null distribution, *x_obs_* is the observed value of mean participation or mean versatility and *I*(⋅) is the indicator function.

#### 2.5.2 What is the relationship between diel structure and network robustness?

To analyse the relationship between diel structure and network robustness we merged the period matrices from each of the 14 networks to obtain aggregated matrices. Then, we simulated random extinctions of plants using the *SimulateExtinctions* function the package NETWORKEXTINCTION v1.0.3 (Ávila-Thieme *et al*. 2023). In these simulations, pollinator species were considered secondarily extinct if they lost more than a specified percentage of their total interaction frequency, referred to as the extinction threshold. Therefore, less abundant and highly specialized visitors are more likely to go secondarily extinct. We tested three thresholds: 60%, 70%, and 80% (function argument IS = 0.4, 0.3, and 0.2). For each network and threshold, simulations were iterated 1,000 times until the network completely collapsed.

Then, we calculated the mean proportion of primary extinctions required to cause the loss half of the species (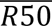) as a measure of robustness. This value was used as the response variable in a generalised linear mixed model (GLMM) with a beta distribution (Ferrari & Cribari-Neto 2004). Fixed effects included mean participation, mean versatility, plant richness (log-transformed), extinction threshold, interaction frequency per network (log-transformed, controlling for sampling intensity), sampling coverage per network, and the number of network layers. We tested a two-way interaction between mean participation and mean versatility to assess potential non-additive effects on robustness. Random effects included a random intercept of the source network nested within site. Because mean participation and the number of network layers were highly collinear (VIF > 5), we included mean participation as the residuals of a linear regression of participation on the number of layers, thereby retaining the component of participation independent of network layers.

#### 2.5.3 How are plant structural traits and floral visual cues associated with the roles plants occupy *across* and *within* diel periods?

The association between plant traits and their roles was assessed at two temporal scales: (1) *across diel periods*, using the 14 networks sampled across multiple diel periods, and (2) *within diel periods*, using the data from all networks. To do so, we implemented three GLMMs with ordered Beta distribution (Kubinec 2023). Fixed effects modelled as linear terms are reported as coefficient estimates (β) with standard errors and Z-statistics, whereas variables modelled as natural splines are reported using Type III Wald χ² tests evaluating the joint significance of all basis functions (see Appendix S4 for model specification details).

##### 2.5.3.1 Effect of plant traits *across diel periods*

To test our predictions on the association of plant traits to the temporal structure of networks, we used the values of plants’ participation (*P_j_*) and versatility (*V_j_*) obtained before. We ran two GLMMs, using participation and versatility as response variables (Appendix S4). Fixed effects included achromatic contrast, colour contrast, plant height, floral complexity, flower abundance, the total interaction frequency per plant, sampling coverage (per network), year, and the number of diel periods (i.e., two or three layers). The random-effects structure included random intercepts for source network nested within site, plant species, and phylogeny. For participation, the model included a zero-inflation component with the total interaction frequency per plant as predictor. For versatility, the model included a dispersion component modelled as a function of the total interaction frequency per plant.

##### 2.5.3.2 Effect of plant traits *within diel periods*

To test for the association of plant traits with the role of plants on specific diel periods, we first calculated centrality for each plant using the period matrices. Centrality quantifies a plant’s structural importance for within-period connectivity. Similar to versatility, to calculate centrality, we first transformed the period matrices into unipartite projections and then we used the PageRank algorithm with the *page_rank* function the package IGRAPH v2.1.1 (Csardi *et al*. 2025; Csardi & Nepusz 2006). In these monolayer matrices, such projections may lead to the deletion of plants that are not connected to the giant component of the network, despite being visited during that period. To avoid missing this information, we added self-loops (with a minimum weight: 1*e* − 10) to the plant nodes so that the random walker of the PageRank algorithm can still teleport to those nodes and assign a centrality value. The values of centrality were then normalized between 0.0 (low importance within the specific period) and 1.0 (high importance within the specific period).

Next, we ran a GLMM using centrality as a response variable. Fixed effects included diel period (“morning”, “afternoon”, or “night”), achromatic contrast, colour contrast, plant height, floral complexity, flower abundance, the total interaction frequency per plant and period, sampling coverage (per period and network), and year. We tested for two-way interactions, between diel period and all plant traits, including floral abundance. The random-effects structure included nested random intercepts for plant species within network within site, a random intercept for plant species, and phylogeny. Additionally, to account for unmeasured, network-specific heterogeneity in night-period conditions (e.g., light conditions) we included a network-level random slope for the night period (see Appendix S4 for justification). Estimated marginal means of trends and contrasts among periods and treatments were estimated using the package EMMEANS v1.11.1 (Lenth 2025), adjusting the results with the Tukey method.

To evaluate whether our model results were robust to the unbalanced diel sampling design across networks, we conducted a series of sensitivity analyses in which each model was refit on progressively restricted subsets of the data (Appendix S5).

## 3 Results

Analysed networks included a total of 123 plant and 508 pollinator species. Overall, we analysed 4853 interaction records involving dipterans (2191), lepidopterans (1427), hymenopterans (1104), coleopterans (110), neuropterans (9), and mecopterans (12) (Fig. 2).

**Figure 2.**
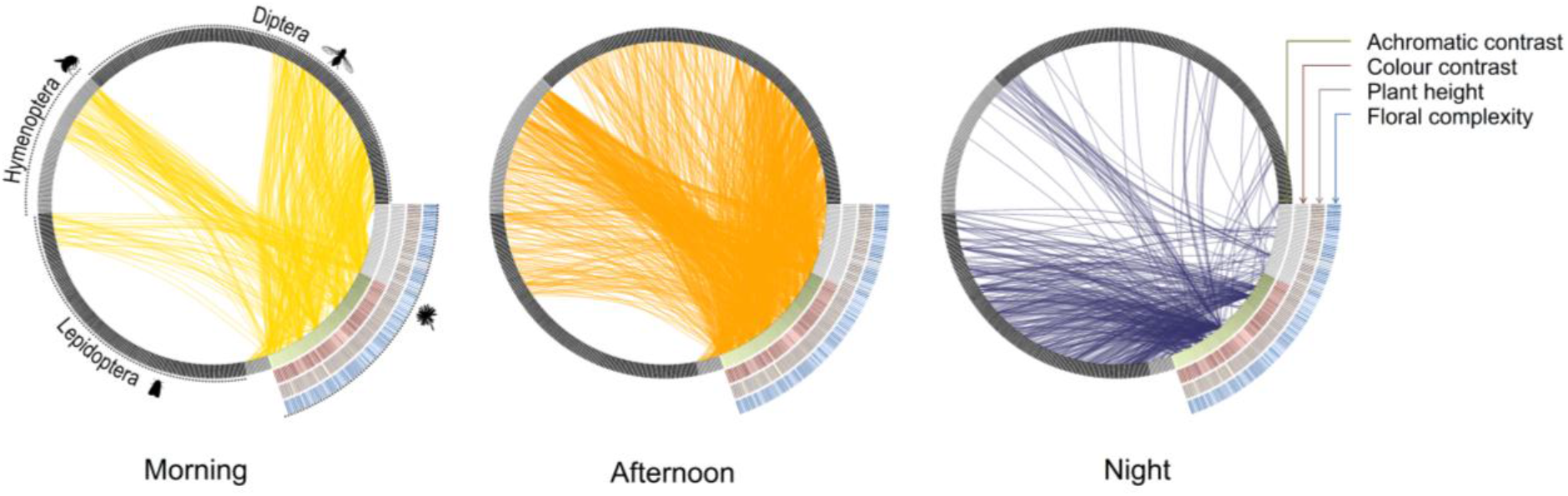
Plant-pollinator interactions across the diel cycle, pooled across networks. Bars represent pollinator (black and dark grey) and plant (coloured) species, and curved lines represent pairwise interactions. Bars adjacent to each plant encode its trait values, with lighter colours indicating higher values and light grey indicating missing data. Species are ordered identically in all panels to allow comparison among diel periods.

### 3.1 The diel structure of plant-pollinator networks exhibits temporal segregation but remains integrated

Mean participation was significantly lower than null expectations in 100%, 93%, and 86% of networks under null model 1, 2, and 3, respectively (Fig. 3A). This suggests that interactions are strongly segregated into diel periods (*P̅* = 0.303 ± 0.147 (mean ± SD); Table S7, Fig. S2, supplementary material). Despite this temporal segregation, mean versatility was significantly higher than null expectations in 93%, 79%, and 71% of networks, respectively (Fig. 3B). These results indicate that networks remain temporally integrated, with certain plant species facilitating cross-period connectivity (*V̅* = 0.205 ± 0.072 (mean ± SD); Table S8, Fig. S2). As observed values significantly differ from all null models, this suggests that the diel structure of these networks arises from non-random preferences in pollinator behaviour, rather than being solely driven by random encounters based on pollinator abundance.

**Figure 3.**
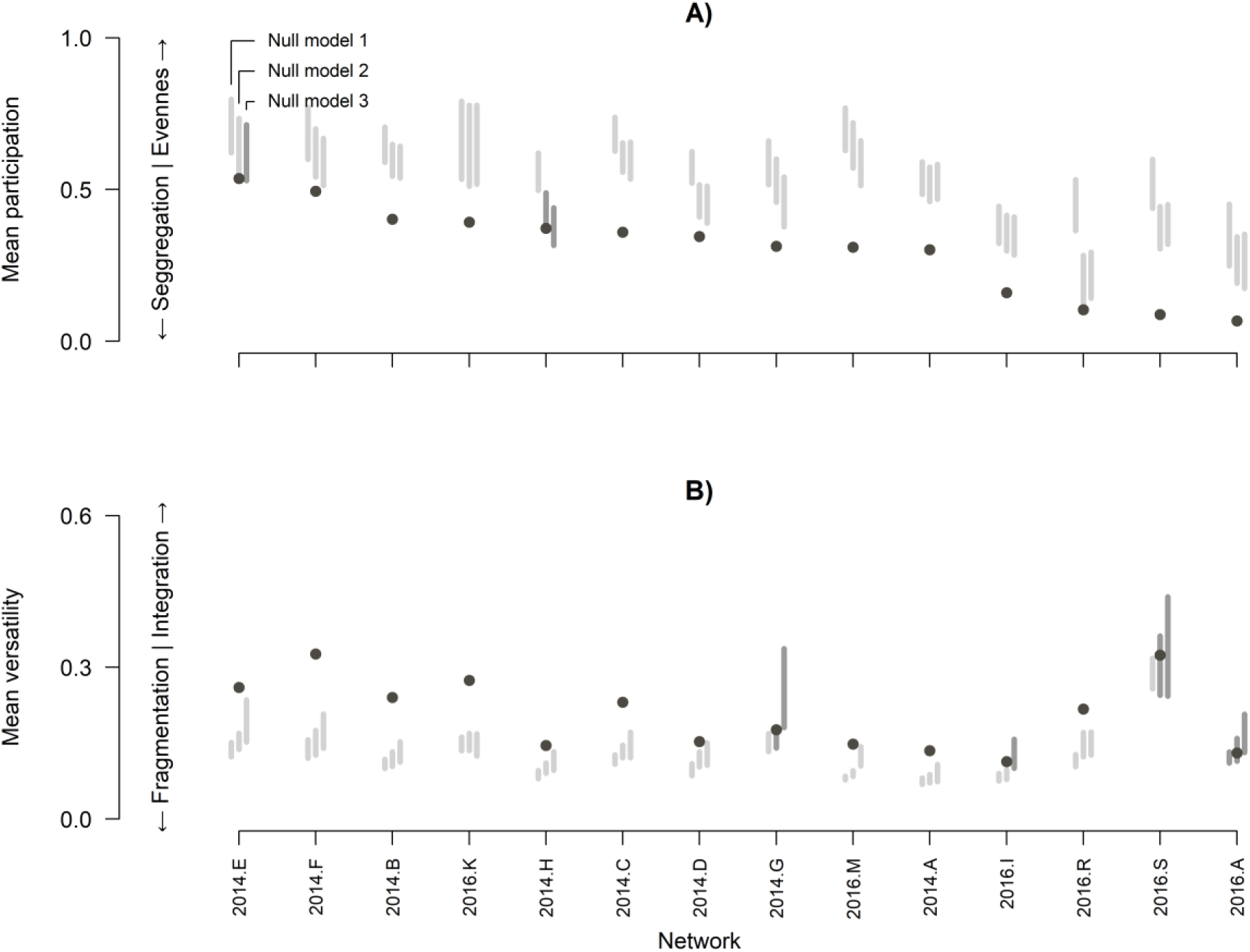
Diel structure of the plant-pollinator networks exhibits both temporal segregation and integration. Observed mean participation (A) and mean versatility (B) of each network (points) are compared against 95% quantile-based confidence intervals (grey lines) from the flexible, layer, and restrictive null models. Darker lines indicate observed values falling within the interval. Networks are ordered by treatment and mean participation, and named by sampling year and site (Table S2).

### 3.2 Robustness is related to the diel structure of plant-pollinator networks

Robustness, estimated as 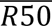, significantly increased with high values of mean versatility (β = 0.057, P < 0.001; Fig. 4), but not with mean participation alone (β = 0.020, P = 0.165). This indicates that diel structures with higher temporal integration tend to be more robust. However, we observed a negative interaction between mean participation and mean versatility (β = -0.058, P < 0.001; Fig 4.), suggesting that high temporal evenness and integration together do not necessarily lead to higher robustness. Robustness also significantly decreased with increasing plant richness (β = -0.023, P = 0.004) but increased with higher sampling coverage (β = 0.047, P < 0.001), the number of network layers (β = 0.080, P < 0.001), and the extinction threshold (β = 0.011, P < 0.001). Interaction frequency per network had no clear effect (β = -0.012, P = 0.189). The sensitivity analysis showed that the direction, magnitude, and significance of the effect all predictors were consistent (Fig. S3, Table S9, Appendix S5).

**Figure 4.**
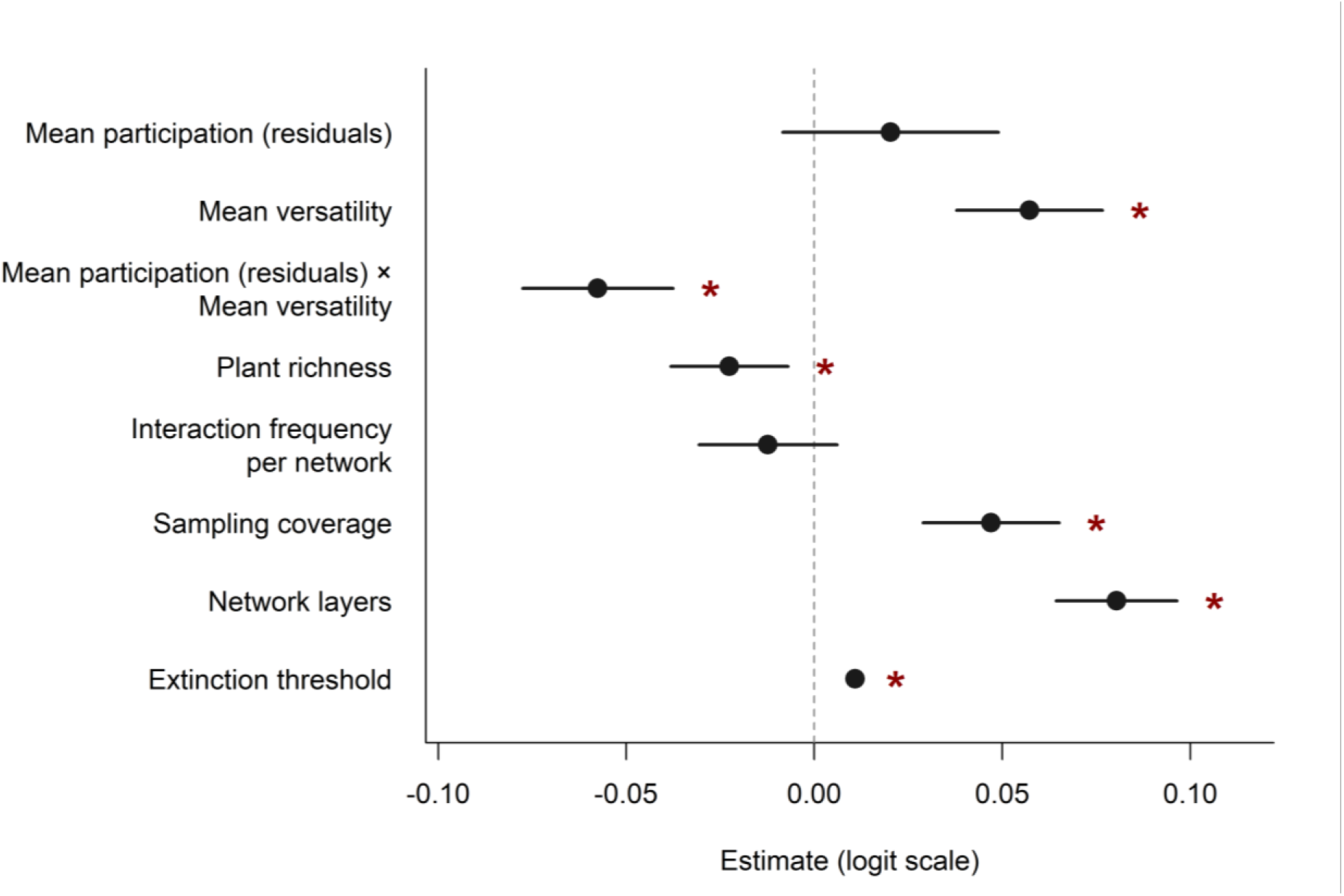
The robustness (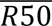) of plant-pollinator networks is related to their diel structure. Points and lines respectively indicate the effect size (β) and the 95% confidence intervals (CI) estimated by the model. Asterisks indicate the cases where confidence intervals do not include 0.

### 3.3 Different plant traits drive the diel structure of plant-pollinator networks *across* and *within* diel periods

*Across diel periods,* structural traits played a key role on diel structure (Fig. 5; Appendix S5). Participation significantly increased with higher plant height (β = 0.288, P = 0.006). Floral complexity, achromatic contrast, colour contrast, flower abundance, and year showed no clear effects (Table S10). Interaction frequency per plant (spline: χ² = 9.998, df = 2, P = 0.007) and the number of network layers had a positive significant effect on participation (β = 0.280, SE = 0.120, P = 0.020), whereas sampling coverage had a significant negative effect (spline: χ² = 9.785, df = 2, P = 0.008). The sensitivity analysis showed that the direction, magnitude, and significance of the effect of plant height were consistent (Figure S4, Table S10, Appendix S5). Therefore, these results suggest that taller plants have their interactions more evenly distributed across periods than smaller ones, which tend to be visited in particular diel periods.

**Figure 5.**
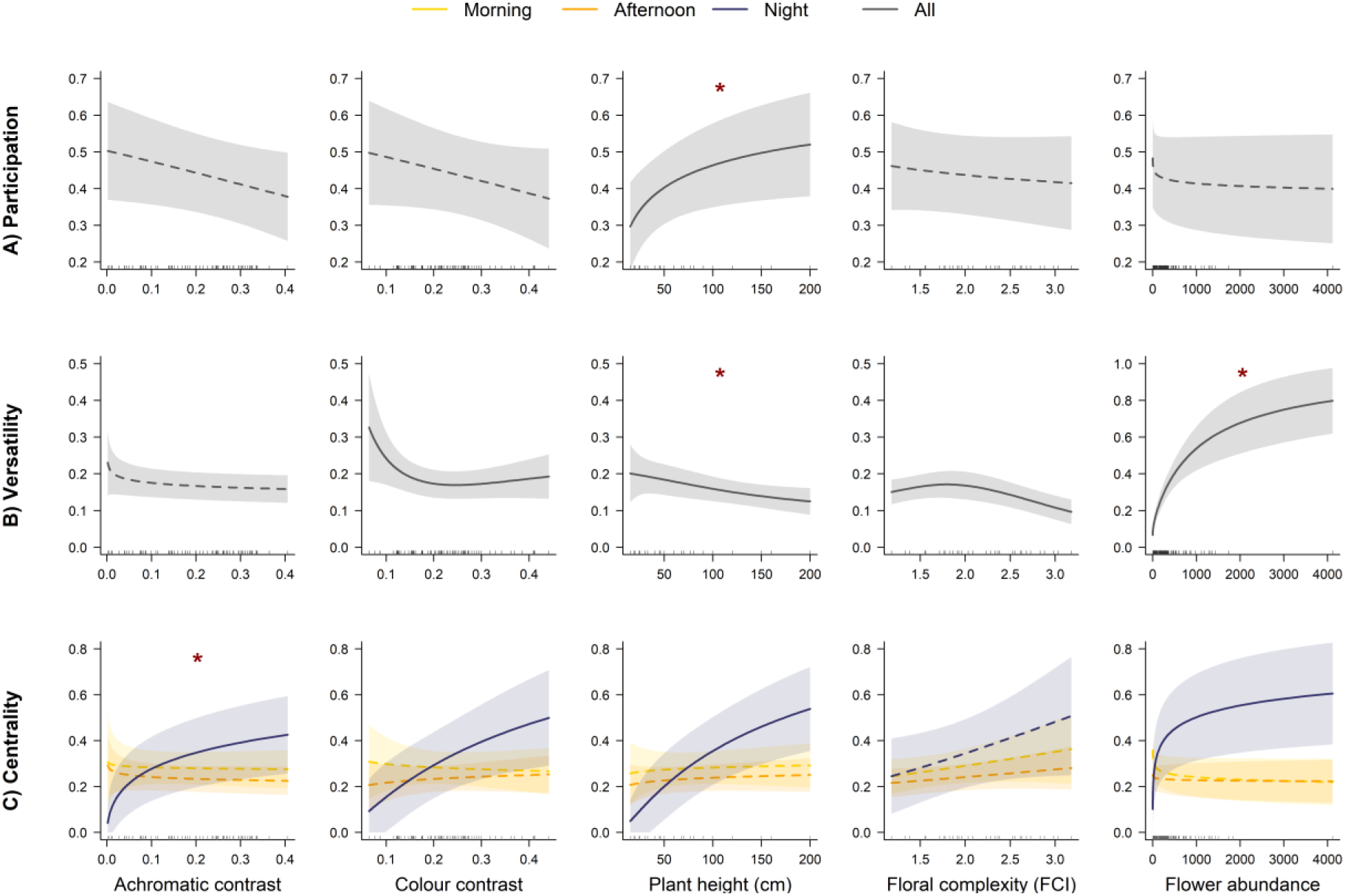
Plant traits are related to the diel structure of plant-pollinator networks at different temporal scales. Results are presented for the GLMMs including participation (A), versatility (B), and centrality (C) as response. Lines and shaded areas indicate the predicted effects and the 95% confidence intervals (CI) estimated by the models with the full dataset. Effects whose P < 0.05 in the models with the full dataset are indicated by solid lines, whereas effects whose P > 0.05 are indicated by dashed lines. The * indicate the most robust predictors based on the sensitivity tests.

Versatility (using the full data set) significantly decreased with higher plant height (spline: χ² = 9.847, df = 2, P = 0.007; Table S11), floral complexity (spline: χ² = 13.480, df = 2, P = 0.001; Fig. 5B; Table S11), and colour contrast (spline: χ² = 11.605, df = 2, P = 0.003; Fig. 5B; Table S11). Conversely, versatility significantly increased with flower abundance (spline: χ² = 151.175, df = 3, P < 0.001; Table S11) and interaction frequency per plant (spline: χ² = 156.135, df = 2, P < 0.001; Fig. 5B; Table S11). Sampling coverage (spline: χ² = 13.281, df = 2, P = 0.001; Fig. 5B; Table S11) and the number of network layers (β = -0.294, SE = 0.101, P = 0.003) had a significantly negative effect. Achromatic contrast and year showed no clear effect on versatility (Table S11). The sensitivity analysis showed that the direction and magnitude of effects were mostly consistent across plant traits (except for colour contrast; Fig. S5). However, among plant traits, only plant height and floral abundance retained significance in the reduced subset (Fig. S5, Table S11, Appendix S5). Therefore, most robust results suggest that short and abundant flowers are the most important for cross-period connectivity.

*Within diel periods*, the balance of influential traits shifted to include visual cues, though structural traits and floral abundance continued to contribute to plant roles at night. These effects are shown by their estimated marginal trends (Est.) and contrasts (Fig. 5C; Table S12—S13). Centrality at night (using the full dataset) was positively influenced by high achromatic contrast (Est. = 0.279, P = 0.001), colour contrast (Est. = 0.233, P = 0.014), plant height (Est. = 0.408, P = 0.001), and flower abundance (Est. = 0.303, P = 0.002); but not during other periods (Fig.5C; Table S12). Floral complexity, year, and sampling coverage had no clear effects on centrality (Table S12). Conversely, centrality increased as interaction frequency per plant and period increased (χ^2^= 173.357, df = 2, P < 0.001; Table S12). Across all sensitivity analyses, the direction and approximate magnitude of trait-by-period effects were consistent across plant traits, but only achromatic contrast showed a consistently significant positive effect in every test (Fig. S6—S7, Appendix S5). All other traits either showed significance only in some subsets or no significant effect at all, highlighting achromatic contrast as the most robust predictor, influencing positively the importance of plants at night for maintaining visitor diversity.

## 4 Discussion

Our study shows that plant-pollinator networks exhibit a non-random diel structure, with interactions temporally segregated into morning, afternoon, and night periods, yet integrated through plants that increase cross-period connectivity. This pattern highlights a functional trade-off in temporal compartmentalisation: temporal segregation likely minimizes competition for pollinators and reduces interspecific pollen transfer (Moreira-Hernández & Muchhala 2019; Phillips *et al*. 2020), whereas temporal integration makes plants more likely to be visited by temporally complementary pollinator groups (Costa *et al*. 2020; Knop *et al*. 2017; Ratoni *et al*. 2024; Zografou *et al*. 2020). The balance between specialising *within* diel periods and generalising *across* them likely contributes to coexistence and the continuity of pollination services (Brosi 2016; Dehling *et al*. 2021). Thus, the moderate compartmentalisation of diel structure may reflect plant strategies that result in coexistence, resource continuity, and overall network functioning.

This diel structure of plant-pollinator networks is closely tied to their robustness: Networks with more temporally even interactions (i.e., networks with many plants whose interactions are uniformly distributed across the diel cycle) and more temporally integrated structures (i.e., networks with many plants which are very well interconnected across periods) showed a higher capacity to sustain function despite species loss. This indicates that the benefits from being visited by temporally complementary pollinator groups may scale from individual plants to the network level (Fontaine *et al*. 2006; Knop *et al*. 2017). However, the negative interaction between mean participation and mean versatility suggests that these benefits have limits. High temporal evenness and temporal integration is an indication of a reduced temporal compartmentalisation, which otherwise may help to contain disturbance spread (Gilarranz *et al*. 2017; Stouffer & Bascompte 2011). Together, these findings show that diel structure is a functional axis of network organisation, where the balance between temporal segregation and integration shapes robustness.

Plant traits determine this diel structure, though their influence varies depending on the temporal scale. *Across diel periods*, structural traits seem to be key drivers, with plant height being the most robust predictor. Taller plants exhibited temporally more evenly distributed interactions (i.e., higher participation), consistent with their greater visibility to potential visitors (Engel & Irwin 2003; Junker *et al*. 2013; Michelot-Antalik *et al*. 2025). In contrast, shorter plants were more important for cross-period connectivity by sharing pollinators with plants from multiple periods (i.e., higher versatility). Depending on the height of surrounding vegetation, low-positioned flowers in grasslands may attract distinct visitor assemblages (Klecka *et al*. 2018). If this vertical stratification effect extends to diel activity patterns, it could help explain why shorter plants seem to play a more important role in connecting a broader range of pollinators across diel periods. Similar positive effects of lower plant height on network roles have been reported at both the individual level (Arroyo-Correa *et al*. 2021) and population level (Aslan *et al*. 2025). This apparent contrast with the participation results is not contradictory: a plant can be visited evenly across diel periods (leading to high participation) without necessarily being visited by a broad range of pollinator taxa on multiple periods (leading to high versatility). Plant abundance also increased versatility, consistent with its known effect on increasing interaction probability in ecological networks (Lázaro *et al*. 2020; Vázquez *et al*. 2007).

*Within diel periods*, the importance of plant traits shifted towards visual cues, with achromatic contrast as the most robust predictor. We expected achromatic cues to be more important at night than during daytime and colour cues being similarly important throughout the diel cycle (van der Kooi & Kelber 2022). Our results partially supported this, as high achromatic contrast was significantly associated with an increased importance of plants for fostering the diversity of pollinators at night (i.e., higher centrality). Plant trait effects on centrality were similar between morning and afternoon, likely because differences are driven by other plant traits, such as scent emission or floral opening (Bloch *et al*. 2017; Fenske & Imaizumi 2016; Fründ *et al*. 2011; Schwarz *et al*. 2021), which were not analysed here. Although our analysis suggests potential effects from other plant traits on versatility and centrality, they remain to be confirmed with more extensive sampling. Overall, these findings support our hypothesis, suggesting that plant-pollinator networks are assembled hierarchically, following both “constant” and “dynamic” rules: *across diel periods*, structural traits set a baseline for interaction potential, whereas *within diel periods*, light conditions determine which traits become most influential.

By incorporating fine-scale temporal dynamics into network ecology, our study provides a mechanistic understanding of the diel structure as a key functional axis of plant-pollinator networks. We show how structural and light-associated traits jointly assemble these networks through a combination of constant and dynamic ecological rules. This framework also highlights that human-driven environmental changes may disproportionately disrupt specific interaction pathways. For example, artificial light at night (ALAN) may affect pathways mediated by achromatic contrast (Briolat *et al*. 2021; Vissio *et al*. 2024), whereas land-use change may primarily disrupt pathways mediated by structural traits (Day Briggs & Anderson 2025; Stefanaki *et al*. 2015). Ultimately, our results demonstrate that diel structure is a critical component of network robustness, shaped by the shifting influence of plant traits across the diel cycle, and that disruptions to these temporal pathways may undermine long-term community persistence.

## Acknowledgments

We thank Natalya Zapata-Mesa and Felix Neff for the interesting discussions about ecological networks and insect ecology. We also thank Melina S. Leite for the suggestions on GLMMs.

## Author contributions

SMB, VG, CF, and EK conceived the idea and methods. SMB and CJvdK compiled functional data and analysed the data. SMB, CJvdK, and EK wrote the first draft of the manuscript. All authors contributed significantly to the final version of the manuscript.

## Data accessibility statement

Associated data and code will be available through Zenodo

## References

Arnold, S.E.J., Faruq, S., Savolainen, V., McOwan, P.W. & Chittka, L. (2010). FReD: The Floral Reflectance Database — A Web Portal for Analyses of Flower Colour. PLOS ONE, 5, e14287.

Arroyo-Correa, B., Bartomeus, I. & Jordano, P. (2021). Individual-based plant–pollinator networks are structured by phenotypic and microsite plant traits. Journal of Ecology, 109, 2832–2844.

Aslan, C.E., Grady, K. & Haubensak, K. (2025). Pollination networks and plant local adaptation: the importance of serving the pollinator community in restoration. Restoration Ecology, 33, e70027.

Ávila-Thieme, M.I., Kusch, E., Corcoran, D., Castillo, S.P., Valdovinos, F.S., Navarrete, S.A., et al. (2023). NetworkExtinction: An R package to simulate extinction propagation and rewiring potential in ecological networks. Methods Ecol. Evol., 14, 1952–1966.

Battiston, F., Nicosia, V. & Latora, V. (2014). Structural measures for multiplex networks. *Phys*. Rev. E, 89, 032804.

Bloch, G., Bar-Shai, N., Cytter, Y. & Green, R. (2017). Time is honey: circadian clocks of bees and flowers and how their interactions may influence ecological communities. Philos. Trans. R. Soc. B, 372, 20160256.

Borges, R.M., Somanathan, H. & Kelber, A. (2016). Patterns and Processes in Nocturnal and Crepuscular Pollination Services. Q. Rev. Biol., 91, 389–418.

Briolat, E.S., Gaston, K.J., Bennie, J., Rosenfeld, E.J. & Troscianko, J. (2021). Artificial nighttime lighting impacts visual ecology links between flowers, pollinators and predators. Nat Commun, 12, 4163.

Brooks, M.E., Kristensen, K., Benthem, K.J. van, Magnusson, A., Berg, C.W., Nielsen, A., et al. (2017). glmmTMB Balances Speed and Flexibility Among Packages for Zero-inflated Generalized Linear Mixed Modeling. R J., 9, 378–400.

Brosi, B.J. (2016). Pollinator specialization: from the individual to the community. New Phytologist, 210, 1190–1194.

CaraDonna, P.J., Burkle, L.A., Schwarz, B., Resasco, J., Knight, T.M., Benadi, G., et al. (2021). Seeing through the static: the temporal dimension of plant–animal mutualistic interactions. Ecol. Lett., 24, 149–161.

CaraDonna, P.J. & Waser, N.M. (2020). Temporal flexibility in the structure of plant– pollinator interaction networks. Oikos, 129, 1369–1380.

Chittka, L. (1992). The colour hexagon: a chromaticity diagram based on photoreceptor excitations as a generalized representation of colour opponency. J. Comp. Physiol. A, 170, 533–543.

Chytrý, M., Řezníčková, M., Novotný, P., Holubová, D., Preislerová, Z., Attorre, F., et al. (2024). FloraVeg.EU — An online database of European vegetation, habitats and flora. Appl. Veg. Sci., 27, e12798.

Costa, J.M., Ramos, J.A., Timóteo, S., da Silva, L.P., Ceia, R.S. & Heleno, R.H. (2020). Species temporal persistence promotes the stability of fruit–frugivore interactions across a 5-year multilayer network. J. Ecol., 108, 1888–1898.

Csardi, G. & Nepusz, T. (2006). The igraph software package for complex network research. InterJournal, Complex Systems, 1695.

Csardi, G., Nepusz, T. & Traag, V. (2025). igraph: Network Analysis and Visualization in R.

Day Briggs, S. & Anderson, J.T. (2025). The effect of global change on the expression and evolution of floral traits. Ann Bot, 135, 9–24.

De Domenico, M. (2022). Multilayer Networks: Analysis and Visualization: Introduction to muxViz with R. Springer International Publishing, Cham.

De Domenico, M., Solé-Ribalta, A., Omodei, E., Gómez, S. & Arenas, A. (2015). Ranking in interconnected multilayer networks reveals versatile nodes. Nat Commun, 6, 6868.

Dehling, D.M., Bender, I.M.A., Blendinger, P.G., Böhning-Gaese, K., Muñoz, M.C., Neuschulz, E.L., et al. (2021). Specialists and generalists fulfil important and complementary functional roles in ecological processes. Funct. Ecol., 35, 1810–1821.

van Doorn, W.G. & Kamdee, C. (2014). Flower opening and closure: an update. J. Exp. Bot., 65, 5749–5757.

van Doorn, W.G. & van Meeteren, U. (2003). Flower opening and closure: a review. J. Exp. Bot., 54, 1801–1812.

Engel, E.C. & Irwin, R.E. (2003). Linking pollinator visitation rate and pollen receipt. American Journal of Botany, 90, 1612–1618.

Fenske, M.P. & Imaizumi, T. (2016). Circadian Rhythms in Floral Scent Emission. Front. Plant Sci., 7.

Ferrari, S. & Cribari-Neto, F. (2004). Beta Regression for Modelling Rates and Proportions. J. Appl. Stat., 31, 799–815.

Fontaine, C., Dajoz, I., Meriguet, J. & Loreau, M. (2006). Functional Diversity of Plant– Pollinator Interaction Webs Enhances the Persistence of Plant Communities. PLoS Biol, 4, e1.

Fründ, J., Dormann, C.F. & Tscharntke, T. (2011). Linné’s floral clock is slow without pollinators – flower closure and plant-pollinator interaction webs. Ecol. Lett.

Gårdman, V., MacDonald, E. & Roslin, T. (2026). Strong diel variation in the activity of insect taxa sampled by Malaise traps. Ecol. Entomol., n/a.

Giavi, S., Blösch, S., Schuster, G. & Knop, E. (2020). Artificial light at night can modify ecosystem functioning beyond the lit area. Sci Rep, 10, 11870.

Giavi, S., Fontaine, C. & Knop, E. (2021). Impact of artificial light at night on diurnal plant-pollinator interactions. Nat Commun, 12, 1690.

Gilarranz, L.J., Rayfield, B., Liñán-Cembrano, G., Bascompte, J. & Gonzalez, A. (2017). Effects of network modularity on the spread of perturbation impact in experimental metapopulations. Science, 357, 199–201.

Giurfa, M., Vorobyev, M., Kevan, P. & Menzel, R. (1996). Detection of coloured stimuli by honeybees: minimum visual angles and receptor specific contrasts. J. Comp. Physiol. A, 178, 699–709.

Grilli, J., Rogers, T. & Allesina, S. (2016). Modularity and stability in ecological communities. Nat. Commun., 7, 12031.

Grognuz, V., Gisler, K. & Knop, E. (2026). Artificial light at night (ALAN) disrupts timing of floral resource availability. Biological Conservation, 314, 111650.

Guimerà, R. & Nunes Amaral, L.A. (2005). Functional cartography of complex metabolic networks. Nature, 433, 895–900.

Hartig, F. (2022). DHARMa: Residual Diagnostics for Hierarchical (Multi-Level / Mixed) Regression Models.

Herrera, C.M. (1990). Daily Patterns of Pollinator Activity, Differential Pollinating Effectiveness, and Floral Resource Availability, in a Summer-Flowering Mediterranean Shrub. Oikos, 58, 277–288.

Hutchinson, M.C., Bramon Mora, B., Pilosof, S., Barner, A.K., Kéfi, S., Thébault, E., et al. (2019). Seeing the forest for the trees: Putting multilayer networks to work for community ecology. Funct. Ecol., 33, 206–217.

Junker, R.R. (2012). Floral Filters: Inviting Mutualists and Screening out Antagonists. Entomol. heute, 24, 21–36.

Junker, R.R., Blüthgen, N., Brehm, T., Binkenstein, J., Paulus, J., Martin Schaefer, H., et al. (2013). Specialization on traits as basis for the niche-breadth of flower visitors and as structuring mechanism of ecological networks. Funct. Ecol., 27, 329–341.

Karolyi, F. (2019). What’s on the Menu: Floral Tissue, Pollen or Nectar? Mouthpart Adaptations of Anthophilous Beetles to Floral Food Sources. In: Insect Mouthparts: Form, Function, Development and Performance (ed. Krenn, H.W.). Springer International Publishing, Cham, pp. 419–442.

Keasar, T. (2020). Patterns of Flower Complexity in Plant Communities. In: Annual Plant Reviews online (ed. Roberts, J.A.). Wiley, pp. 643–660.

Kelber, A., Vorobyev, M. & Osorio, D. (2003). Animal colour vision — behavioural tests and physiological concepts. Biol. Rev., 78, 81–118.

Kendall, L. & Nicholson, C.C. (2025). Pollination Across the Diel Cycle: A Global Meta-Analysis. Ecol. Lett., 28, e70036.

Klecka, J., Hadrava, J. & Koloušková, P. (2018). Vertical stratification of plant–pollinator interactions in a temperate grassland. PeerJ, 6, e4998.

Knop, E., Gerpe, C., Ryser, R., Hofmann, F., Menz, M.H.M., Trösch, S., et al. (2018). Rush hours in flower visitors over a day–night cycle. Insect Conserv. Divers., 11, 267–275.

Knop, E., Zoller, L., Ryser, R., Gerpe, C., Hörler, M. & Fontaine, C. (2017). Artificial light at night as a new threat to pollination. Nature, 548, 206–209.

van der Kooi, C.J., Dyer, A.G., Kevan, P.G. & Lunau, K. (2019). Functional significance of the optical properties of flowers for visual signalling. Ann. Bot., 123, 263–276.

van der Kooi, C.J., Elzenga, J.T.M., Staal, M. & Stavenga, D.G. (2016a). How to colour a flower: on the optical principles of flower coloration. Proc. R. Soc. B, 283, 20160429.

van der Kooi, C.J. & Kelber, A. (2022). Achromatic Cues Are Important for Flower Visibility to Hawkmoths and Other Insects. Front. Ecol. Evol., 10.

van der Kooi, C.J., Pen, I., Staal, M., Stavenga, D.G. & Elzenga, J.T.M. (2016b). Competition for pollinators and intra-communal spectral dissimilarity of flowers. Plant Biol J, 18, 56–62.

van der Kooi, C.J. & Spaethe, J. (2025). Flower colour contrast, ‘spectral purity’ and a red herring. Plant Biol., 27, 189–194.

Krenn, H.W. (2019). Form and Function of Insect Mouthparts. In: Insect Mouthparts: Form, Function, Development and Performance (ed. Krenn, H.W.). Springer International Publishing, Cham, pp. 9–46.

Krenn, H.W., Plant, J.D. & Szucsich, N.U. (2005). Mouthparts of flower-visiting insects. Arthropod Struct. Dev., 34, 1–40.

Krishna, S. & Keasar, T. (2018). Morphological Complexity as a Floral Signal: From Perception by Insect Pollinators to Co-Evolutionary Implications. Int. J. Mol. Sci., 19, 1681.

Kubinec, R. (2023). Ordered Beta Regression: A Parsimonious, Well-Fitting Model for Continuous Data with Lower and Upper Bounds. Polit. Anal., 31, 519–536.

Lauber, K., Wagner, G. & Gygax, A. (2018). Flora Helvetica – Illustrated Flora of Switzerland. 7th edn. Haupt Verlag.

Lázaro, A., Gómez-Martínez, C., Alomar, D., González-Estévez, M.A. & Traveset, A. (2020). Linking species-level network metrics to flower traits and plant fitness. J. Ecol., 108, 1287–1298.

Lenth, R.V. (2025). emmeans: Estimated Marginal Means, aka Least-Squares Means.

Lüdecke, D., Ben-Shachar, M.S., Patil, I., Waggoner, P. & Makowski, D. (2021).performance: An R Package for Assessment, Comparison and Testing of Statistical Models. J. Open Source Softw., 6, 3139.

Maia, K.P., Vaughan, I.P. & Memmott, J. (2019). Plant species roles in pollination networks: an experimental approach. Oikos, 128, 1446–1457.

Martín González, A.M., Dalsgaard, B. & Olesen, J.M. (2010). Centrality measures and the importance of generalist species in pollination networks. Ecol. Complex., 7, 36–43.

May, R.M. (1972). Will a Large Complex System be Stable? Nature, 238, 413–414.

Michelot-Antalik, A., Langlois, A., de Bello, F., Desaegher, J., Genty, L., Goulnik, J., et al. (2025). Handbook of protocols for standardized measurements of floral traits for pollinators in temperate communities. Methods Ecol. Evol., 16, 988–1001.

Moreira-Hernández, J.I. & Muchhala, N. (2019). Importance of Pollinator-Mediated Interspecific Pollen Transfer for Angiosperm Evolution. Annu. Rev. Ecol. Evol. Syst., 50, 191–217.

Olesen, J.M., Bascompte, J., Dupont, Y.L. & Jordano, P. (2007). The modularity of pollination networks. Proc. Natl. Acad. Sci. U.S.A., 104, 19891–19896.

Phillips, R.D., Peakall, R., Van Der Niet, T. & Johnson, S.D. (2020). Niche Perspectives on Plant–Pollinator Interactions. Trends Plant. Sci., 25, 779–793.

Pilosof, S., Porter, M.A., Pascual, M. & Kéfi, S. (2017). The multilayer nature of ecological networks. Nat Ecol Evol, 1, 1–9.

Ratoni, B., Cruz, C.P., Novais, S., Rodríguez-Morales, D., Neves, F.S., Ayala, R., et al. (2024). Temporal decay of similarity in bee-plant relationships throughout the day. Oecologia, 207, 1–12.

Roemer, R.B., Irene Terry, L., Booth, D.T. & Walter, G.H. (2022). Insights from an ancient gymnosperm lineage: ambient temperature and light and the timing of thermogenesis in cycad cones. Am. J. Bot., 109, 151–165.

Schwarz, B., Dormann, C.F., Vázquez, D.P. & Fründ, J. (2021). Within-day dynamics of plant–pollinator networks are dominated by early flower closure: an experimental test of network plasticity. Oecologia, 196, 781–794.

Schwarz, B., Vázquez, D.P., CaraDonna, P.J., Knight, T.M., Benadi, G., Dormann, C.F., et al. (2020). Temporal scale-dependence of plant–pollinator networks. Oikos, 129, 1289–1302.

Souza, C.S., Oliveira, P.E., Rosa, B.B. & Maruyama, P.K. (2022). Integrating nocturnal and diurnal interactions in a Neotropical pollination network. J. Ecol., 110, 2145–2155.

Spaethe, J., Tautz, J. & Chittka, L. (2001). Visual constraints in foraging bumblebees: Flower size and color affect search time and flight behavior. Proc. Natl. Acad. Sci. U.S.A., 98, 3898–3903.

Stefanaki, A., Kantsa, A., Tscheulin, T., Charitonidou, M. & Petanidou, T. (2015). Lessons from Red Data Books: Plant Vulnerability Increases with Floral Complexity. PLOS ONE, 10, e0138414.

Stouffer, D.B. & Bascompte, J. (2011). Compartmentalization increases food-web persistence. Proc. Natl. Acad. Sci. U.S.A., 108, 3648–3652.

Teng, Y., Villalobos, S., Vamosi, J.C., Wang, X.-F. & Gong, Y.-B. (2024). Diurnal versus nocturnal pollination in a subalpine wetland: From network structure to plant reproduction. Glob. Ecol. Conserv., 49, e02798.

Thieurmel, B. & Elmarhraoui, A. (2025). suncalc: Compute Sun Position, Sunlight Phases, Moon Position and Lunar Phase.

Vázquez, D.P., Melián, C.J., Williams, N.M., Blüthgen, N., Krasnov, B.R. & Poulin, R. (2007). Species abundance and asymmetric interaction strength in ecological networks. Oikos, 116, 1120–1127.

Vissio, C., Drewniak, E.M., Cocucci, A.A., Moré, M., Benitez-Vieyra, S., Giaquinta, A., et al. (2024). Artificial light changes visual perception by pollinators in a hawkmoth-plant interaction system. Urban Ecosyst., 27, 1235–1249.

Xu, X., Ren, Z.-X., Trunschke, J., Kuppler, J., Zhao, Y.-H., Knop, E., et al. (2021). Bimodal activity of diurnal flower visitation at high elevation. Ecol. Evol., 11, 13487–13500.

Zografou, K., Swartz, M.T., Tilden, V.P., McKinney, E.N., Eckenrode, J.A. & Sewall, B.J. (2020). Stable generalist species anchor a dynamic pollination network. Ecosphere, 11, e03225.

Zoller, L., Bennett, J.M. & Knight, T.M. (2020). Diel-scale temporal dynamics in the abundance and composition of pollinators in the Arctic summer. Sci Rep, 10, 21187.

