## Supplementary material for "The diel structure and robustness of plant-pollinator networks are driven by the shifting influences of plant traits"

---

### Appendix S1: Further details on study site and interaction sampling

The study was conducted in the Swiss Prealps, a biogeographical and geological unit characterized predominantly by sedimentary rocks and situated in the lower frontal ranges north of the central Alps (1, 2). Study sites were located at elevations ranging approximately from 650 to 1000 m a.s.l.. The mean (± SD) distance between sites varied among years, measuring 2.58 ± 1.60 km in 2014, 3.11 ± 2.45 km in 2015, 12.91 ± 7.73 km in 2016, and 10.70 ± 7.20 km in 2017. All sites were comparable in vegetation, with *Cirsium oleraceum* (Asteraceae) usually being the most abundant.

Sampling was conducted during spring and summer, spanning May-September in 2014, June-September in 2015 and 2016, and June-August in 2017. At each site, flower-visitor interactions were sampled along a 100 m transect (reduced to 50 m in 2014). All flower visitors contacting the reproductive organs of flowers within 1 m on either side of the transect were collected using a hand net while walking at a constant pace (3). Each individual was captured separately and immediately placed into an individual vial to prevent cross-contamination. For every captured visitor, the associated plant species was recorded. Specimens were subsequently frozen, pinned, and identified to species level whenever possible.

Sampling was conducted during morning (from ~08:00), afternoon (from ~13:30), and night (starting ~30 min after astronomical sunset) periods, each lasting up to four hours. Nocturnal sampling was typically paired with diurnal sampling (weather permitting), with multiple periods surveyed on the same day or on adjacent days. Sampling was restricted to suitable weather conditions (i.e. no strong wind or rain, generally sunny during the day). Within each sampling diel period, transects were surveyed at 30-minute intervals. Sites were visited repeatedly throughout the sampling period. In 2014, sites were visited six times; in 2015, five times (one site four times); and in 2016-2017, six to seven times.

Sampling coverage of plants was (mean ± SD) 0.999 ± 0.002, for pollinators it was 0.830 ± 0.071, whereas for interactions 0.667 ± 0.132 (Table S2). Sampling coverage of pollinators per period was 0.685 ± 0.176 for morning, 0.806 ± 0.062 for afternoon, and 0.725 ± 0.164 for night. Sampling coverage of interactions per period was (mean ± SD) 0.492 ± 190 for morning, 0.579 ± 0.123 for afternoon, and 0.610 ± 0.206 for night.

### Appendix S2: Weights of floral traits for the calculation of the floral complexity index.

Following Stefanaki et al. (4), the traits used to calculate the Floral Complexity Index (FCI) are presented below with their corresponding "complexity weights" in parenthesis:

a) Floral shape. Flowers were assigned into nine different categories according to their functional shape: bell (1.00), brush (1.05), disk (0.30), tube (0.98), disk-tube (0.83), funnel (0.80), flag (1.28), gullet (1.13), or head (0.68).

b) Floral depth. Depending on the length of the corolla tube, plant species were assigned to three categories: low-depth flowers (0.20), for corolla tube length < 4 mm; medium-depth flowers (0.40), for corolla tube length between 4 and 10 mm; and high-depth flowers (0.60), for corolla tube length > 10 mm.

c) Floral symmetry. Plant species were assigned a category, depending on the number of floral symmetry axes, as radial (0.23), for those with several axes, and bilateral (0.50), for those with a single axis.

d) Corolla segmentation. Depending on the corolla, they were classified as sympetaly (0.50), chorypetaly (0.40), and semichoripetaly (0.30).

e) Functional reproductive unit. Depending on the type of inflorescence, plant species were assigned to three possible levels: single/few flowers (0.15), flat or spherical segregations (0.15), and cylindrical segregations (0.30).

### Appendix S3: Preparing networks for the calculation of versatility

For the calculation of versatility, we had first to build temporal multilayer bipartite networks using the *create multilayer network* function of the EMLN package v1.0.2 (5). Each network included four components as described below:

(a) Layers representing diel periods: morning ($\propto$), afternoon ($\beta$), and night ($\gamma$).

(b) Two sets of nodes representing pollinator and plant species in all layers.

(c) Intralayer directed weighted links connecting plant $j$ with pollinator $i$, and vice versa, in layer $l = [\propto, \beta, \gamma]$, defined as $w_{ji}^{l} = w_{ij}^{l} = f_{ji}/T^{l}$, where $f_{ji}$ denotes the number of visits of $i$ to $j$, and $T^{l}$ denotes the total interaction frequency in layer $l$.

(d) Interlayer directed weighted links connecting each pollinator and plant species with itself, where the direction of the links point towards the "future" layer. Although the three different layers that were analysed depict a temporal component of plant-pollinator interactions, they occur in a cyclical manner (i.e., morning → afternoon → night → morning), so we used interlayer directed links to capture that periodicity. For each pollinator, its interlayer link weights were calculated as the ratio between the number of visits in the "next" layer in relation to the number of visits in the "current" layer and divided by the total number of visits of the network. For instance, the interlayer directed link weight of pollinator $i$ from the morning ($\propto$) to afternoon ($\beta$) layers was calculated as $w_{i}^{\propto\beta} = t_{i}^{\beta}/(t_{i}^{\propto}T)$, where $t_{i}^{\propto}$ and $t_{i}^{\beta}$ are the number of visits of pollinator $i$ to all plants in layers $\propto$ and $\beta$, respectively, and $T$ is the total interaction frequency of the network. On the other hand, for plants, interlayer link weights were calculated as the relative flower abundance of each species during the whole study period, which means that for each plant species in a network its interlayer link weights were the same among layers. For instance, the interlayer link weight of plant $j$ between all morning ($\propto$), afternoon ($\beta$), and night ($\gamma$) layers was calculated as $w_{j}^{\propto\beta\gamma} = a_{j}/A$, where $a_{j}$ is the flower abundance of plant $j$, and $A$ is the summed flower abundance of all plant species in the multilayer network. This way, both intra- and interlayer link weights range between 0 and 1, reducing potential bias due to different weight scales (6).

Then, these multilayer networks were transformed into unipartite projections, where plants within periods were linked depending on whether they share a pollinator (7, 8). These projections were obtained using the *projecting tm* function of the package TNET v3.0.16 (9) with the Newman (2001) method, which considers interaction weights based on the shared pollinators and the assemblage size. Interlayer links for plants do not change, conserving the temporal relationship among layers. Interlayer links for pollinators are not considered in this projection. Despite projecting periods into unipartite projections may lead to the deletion of plants that are not connected to the giant component in monolayer networks, we did not have such problem with our multilayer projections, as these plants are conserved in our analysis thanks to their interlayer links.

### Appendix S4: Details on structure of GLMMs

All three models (i.e., for participation, versatility, and centrality) were fitted as generalised linear mixed models with ordered beta distributions. The ordered beta distribution was chosen because the three response variables are bound between zero and one and may include observations at both boundaries, which standard beta regression cannot accommodate. All continuous predictors were centred and standardised (mean = 0, SD = 1) prior to analysis to facilitate comparison of effect sizes across variables measured on different scales.

*Variable transformations*

The following variables were log-transformed prior to standardisation to reduce right-skewness and improve linearity of relationships with the response: plant height (all three models), floral abundance (all three models), achromatic contrast (versatility and centrality models), colour contrast (versatility and centrality models), floral complexity (participation model only), interaction frequency (participation and centrality models), and sampling coverage (centrality model only). Network layers and year were not log-transformed in any model. The choice of transformation for each variable in each model was guided by inspection of the raw distributions and residual diagnostics within each dataset. All continuous predictors were centred and standardised (mean = 0, SD = 1) after any log-transformation to facilitate comparison of effect sizes across variables measured on different scales.

*Use of natural splines*

Predictors were modelled as linear terms by default. Natural splines were introduced only when there was evidence that a nonlinear specification improved model fit, assessed through a combination of DHARMa residual diagnostics (visual inspection of quantile residuals and the quantile test), comparison of AIC between linear and spline specifications, Type III Wald $\chi^2$ tests, and likelihood ratio tests confirming that the additional spline parameters contributed significantly to model fit. The number of degrees of freedom for each spline (2 or 3) was selected as the minimum necessary to capture the observed nonlinearity without overfitting, guided by the same diagnostic criteria.

*Participation model*

In the participation model (logit link), all floral traits (achromatic contrast, colour contrast, floral complexity, plant height, floral abundance), interaction frequency, network layers (2 or 3 diel periods), and year were modelled as linear terms. Sampling coverage was modelled as a natural spline with two degrees of freedom. A zero-inflation component with interaction frequency as a predictor was included to account for the excess of zero values in participation at low sampling intensity, where insufficient observation time prevents the detection of interactions and participation is consequently estimated as zero. The random-effects structure included random intercepts for source network nested within site, plant species, and phylogeny. This model was coded as:

```
Participation ~

Achromatic_contrast + Colour_contrast + Floral_complexity + Plant_height + Floral_abundance + Interaction_frequency + ns(Sampling_coverage, 2) + Network_layers + Year + (1 | Plant_species) + (1 | Site/Source_network) + propto(0 + Plant_species | phylo_dummy, A)

Zero-inflation: ~ interaction frequency
```

where `phylo_dummy` is a single-level grouping factor that assigns all observations to a common group (a technical requirement of glmmTMB for specifying custom covariance structures across species), and `A` is a phylogenetic correlation matrix derived from the plant phylogeny.

*Versatility model*

In the versatility model (logit link), colour contrast, floral complexity, plant height, floral abundance, interaction frequency, and sampling coverage were modelled as natural splines with two degrees of freedom each. Achromatic contrast, network layers (2 or 3 diel periods), and year were modelled as linear terms. The dispersion parameter was modelled as a function of interaction frequency (natural spline with two degrees of freedom) and floral abundance (natural spline with three degrees of freedom) to account for heteroscedasticity associated with variation in sampling intensity and abundance. The random-effects structure included random intercepts for source network nested within site, plant species, and phylogeny. This model was coded as:

```
Versatility ~

Achromatic_contrast + ns(Colour_contrast, 2) + ns(Floral_complexity, 2) + ns(Plant_height, 2) + ns(Floral_abundance, 3) + ns(Interaction_frequency, 2) + Sampling_coverage + Network_layers + Year + (1 | Plant_species) + (1 | Site/Source_network) + propto(0 + Plant_species | phylo_dummy, A)

Dispersion: ~ ns(Interaction_frequency, 2) + ns(Floral_abundance, 3)
```

*Zero-inflation and dispersion components*

The zero-inflation component in the participation model and the dispersion component in the versatility model were introduced after inspection of DHARMa quantile residuals. Diagnostics revealed systematic patterns indicative of excess zeros and heteroscedasticity, respectively; their inclusion resolved these diagnostic issues and improved model fit.

*Centrality model*

In the centrality model (probit link), interaction frequency (per plant and period) was modelled as a natural spline with two degrees of freedom. All five floral traits (achromatic contrast, chromatic contrast, floral complexity, plant height, floral abundance) were modelled as linear terms interacting with diel period (i.e., morning, afternoon, night), allowing the relationship between each trait and centrality to vary across diel periods. Sampling coverage and year were modelled as linear terms. The centrality model used a probit link rather than the logit link used for participation and versatility, as DHARMa quantile residual diagnostics under the logit link showed systematic departures that were resolved under the probit specification.

Random effect terms were added according to the following reasoning:

- *Random effect term:* nested random intercepts for plant species within network within site.
  *Reasoning:* to account for the hierarchical structure of the data, where repeated observations of the same plant species across diel periods are nested within networks, which are in turn nested within sites.
  *Code as in glmmTMB:* `(1| Site/Source_network/Plant_species)`

- *Random effect term:* a crossed random intercept for plant species.
  *Reasoning:* to account for species variation that cannot be attributed to phylogeny.
  *Code as in glmmTMB:* `(1| Plant Species)`

- *Random effect term:* phylogeny.
  *Reasoning:* To account for phylogenetic non-independence among plant species.
  *Code as in glmmTMB:* `(0 + Plant species | phylo_dummy, A)`

- *Random effect term:* network-level random slope for the night period.
  *Reasoning:* Unlike morning and afternoon sampling, which occurred under direct sunlight and therefore experienced comparatively "standardized" ambient illumination across networks, night-period sampling occurred in the absence of a dominant light source. Night-period visits were distributed across a wide range of lunar phases in a non-systematic manner, and ambient light conditions at night may have varied further according to cloud cover, canopy structure, or other local site characteristics.
  *Code as in glmmTMB:* `(0 + Night_ind | Site/Source_network)`, where `Night_ind` is a binary indicator coded 1 for night-period observations and 0 otherwise, allowing each network's deviation in night-period activity to be estimated independently rather than assumed constant across networks.

This model was coded as:

```
Centrality ~ Diel_period × (Achromatic_contrast + Colour_contrast + Floral_complexity +
Plant_height + Floral_abundance) + ns(Interaction_frequency, 2) + Sampling_coverage + Year
+ (1 | Site/Source_network/Plant_species) + (1 | Plant_species) + (0 + Night_ind |
Source_network)+ propto(0 + Plant_species | phylo_dummy, A)
```

### Appendix S5: Sensitivity tests

To evaluate whether our models' results were robust to the unbalanced diel sampling design across networks, we conducted a series of sensitivity analyses in which each model was refit on progressively restricted subsets of the data.

*Robustness*

Starting from the full dataset of networks sampled during multiple diel periods (42 observations, 14 networks), we removed all networks sampled only during the afternoon and night (yielding 33 observations, 11 networks). For each dataset (the full dataset and the subset), the models were refit with an identical structure, and estimates, standard errors and associated statistics were extracted. The effect of all predictor variables yielded similar results when using the full dataset and the subset (Table S9, Fig. S3).

*Participation and Versatility*

Starting from the full dataset of networks sampled during multiple diel periods (245 observations, 14 networks), we removed all networks sampled only during the afternoon and night (yielding 347 observations, 11 networks). For each dataset (the full dataset and the subset), the models were refit with an identical structure, and estimates, standard errors and associated statistics were extracted.

For participation, the effect of all predictor variables yielded similar results when using the full dataset and the subset (Table S10, Figure S4), with plant height showing a significant positive effect on participation (Figure S4D). For versatility, most predictor variables also yielded similar results when using the full dataset and the subset, but among plant traits only the effect of plant height was significant in all cases (Table S11, Figure S5).

*Centrality*

Starting from the full dataset (448 observations, 22 networks), we first removed all networks sampled only at night (yielding 407 observations, 14 networks). From this reduced dataset, we generated two further subsets: one excluding morning observations entirely, retaining only networks with afternoon and night data (293 observations, 14 networks), and one additionally excluding all networks sampled only during afternoon and night, retaining only networks with complete diel coverage across all three periods (347 observations, 11 networks). Note that centrality is a layer-level metric, therefore, removing the morning observations from a network does not influence the values observed in other diel periods. For each of the four datasets (the full dataset and three sensitivity subsets), the model was refit with an identical structure, and estimated marginal trends and their 95% confidence intervals were extracted for each period and trait.

Across all four sensitivity tests, the direction and approximate magnitude of trait-by-period effects remained consistent for most traits (Table S12, Figure S6). Achromatic contrast showed significant positive night-period trends (95% CI excluding zero) in the full dataset and in all three sensitivity subsets (Figure S5A). The positive effect of colour contrast at night was only significant with the full data set and the subset of networks when the night-only networks were removed. Plant height and floral abundance showed significant positive night-period trends, except for the subset of networks sampled in all three periods (Figure S5B, D, E). Floral complexity showed no significant night-period effect in any of the four datasets (Figure S6C). Sampling coverage per network showed a consistent negative (non-significant to marginal) trend in the full dataset and two of the three sensitivity subsets, but reversed to a significant positive trend in the subset excluding both night-only networks and morning observations, suggesting this particular covariate's estimated relationship with the response is sensitive to the specific data subset retained (Figure S6F). Year and interaction frequency per plant and period, the remaining period-invariant covariates, showed stable estimates across all four datasets (Figure S6G-H). Together, these results indicate that the focal trait-by-night effects are not driven by the unequal distribution of diel sampling across networks and remain detectable even when the analysis is restricted to the subset of networks with complete morning, afternoon, and night coverage.

Such robustness within our results was also observed in the pairwise contrasts between diel periods for the association between floral traits and plant centrality (Figure S7). The effect of achromatic contrast on plant centrality was significantly higher at night than during the other periods (Table S13), except when we removed the night-only and the afternoon-night-only networks. In this last specific case, the effect of achromatic contrast at night was only significantly higher at night than during the afternoon (Figure S7A).

### Figures

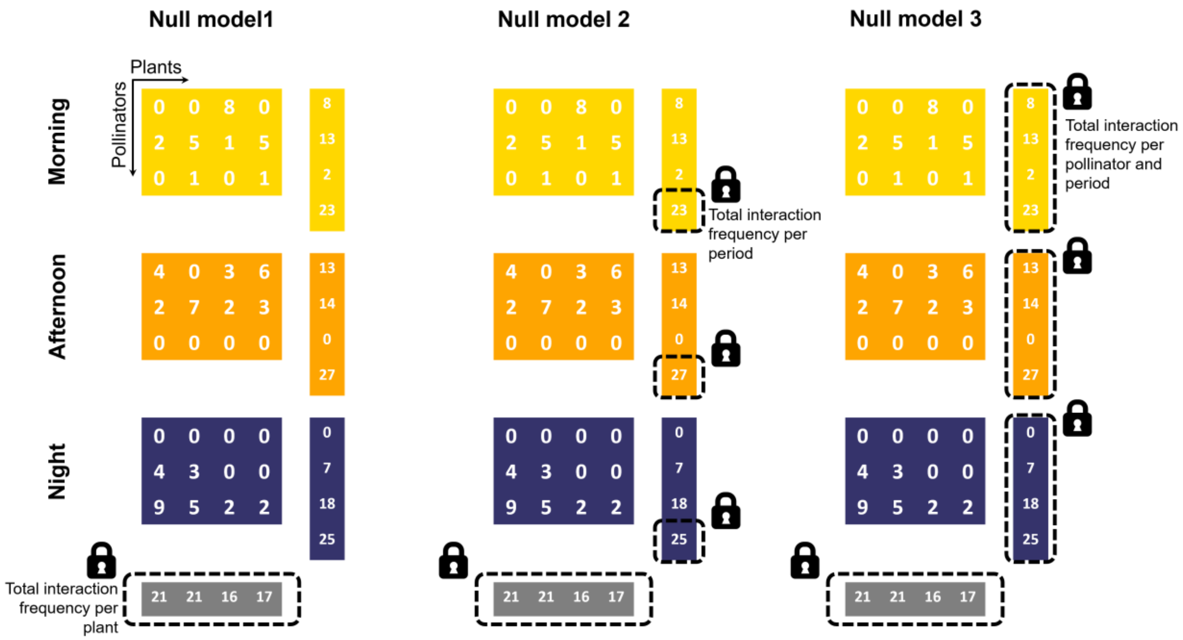

**Figure S1.** The temporal null models. Each model constrained a different part of the observed multilayer networks, which is indicated by the dashed rectangles and the black locks. By comparing the observed mean participation and mean versatility against distributions generated from these three null models, we determined whether temporal structure emerged beyond constraints of (1) overall plant interaction frequency, (2) period-specific interaction frequency, or (3) period-specific pollinator interaction frequency.

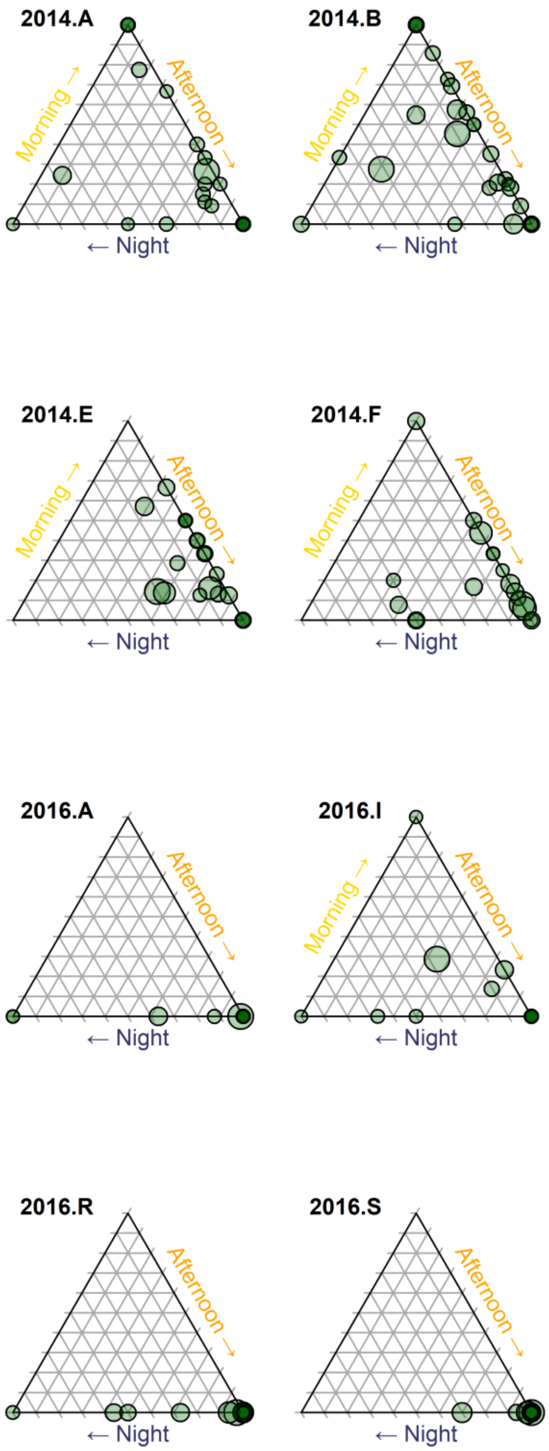

**Figure S2.** Interaction distribution of the 14 networks with data from multiple diel periods. Plant species from each network (points) are arranged in ternary plots based on the proportion of their interactions occurring at each period. This means that plants closer to the centre have higher participation ($P_j$) than those closer to the corners. Point size is proportional to the versatility ($V_j$) of each plant.

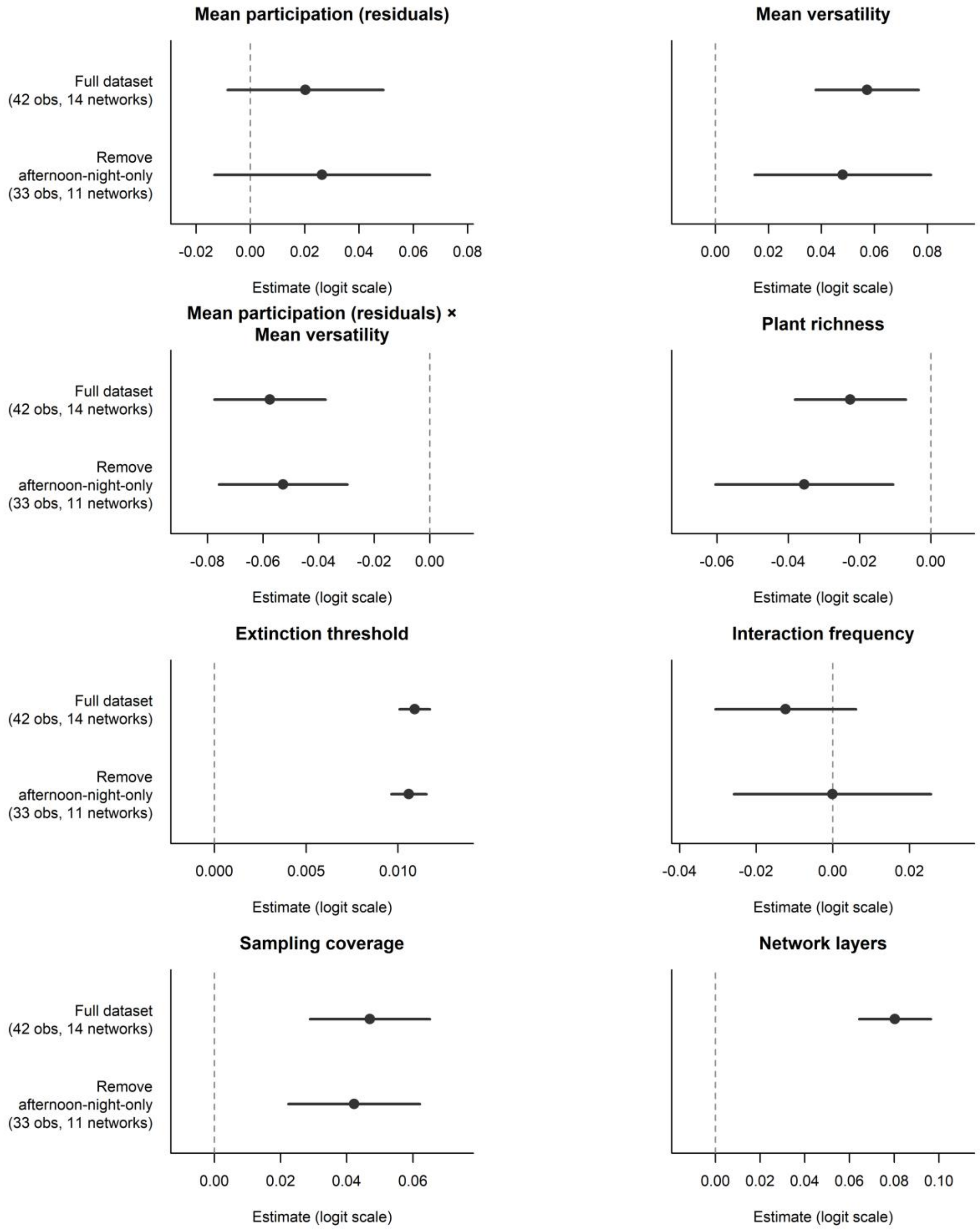

**Figure S3.** Results from the sensitivity test on the association between diel structure and robustness. Rows from top to bottom show the estimated fixed effects for the model with the full set of networks and after removing networks sampled during afternoon and night only. Estimates are indicated by the dots and lines depict 95% confidence intervals.

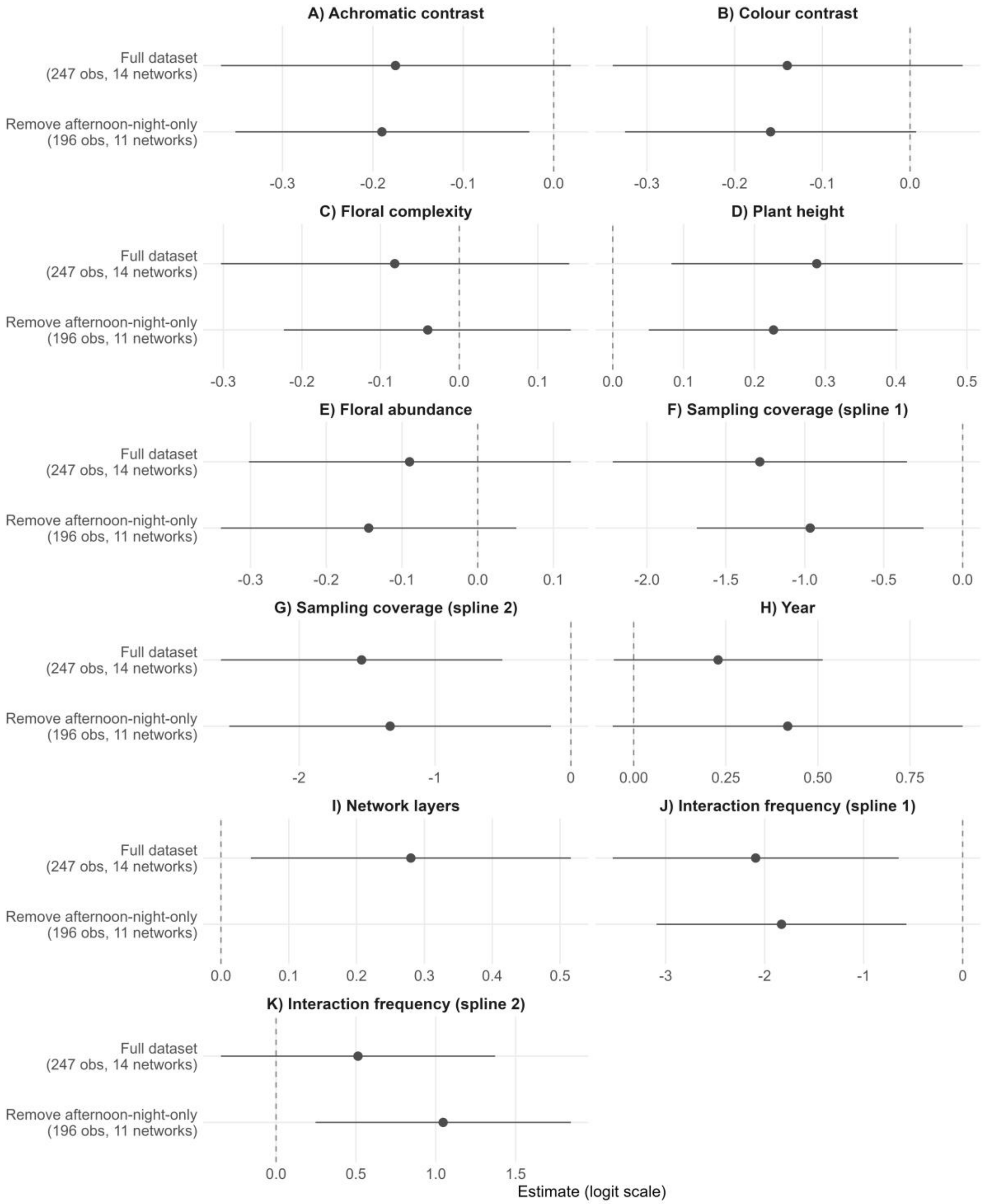

**Figure S4.** Results from the sensitivity test on the association between floral traits and participation. Rows from top to bottom show the estimated fixed effects for the model with the full set of networks and after removing networks sampled during afternoon and night only. Estimates are indicated by the dots and lines depict 95% confidence intervals. A) Achromatic contrast, B) colour contrast, C) floral complexity, D) plant height, E) Floral abundance, F-G) Sampling coverage, H) year, I) layers, J-K) interaction frequency per plant.

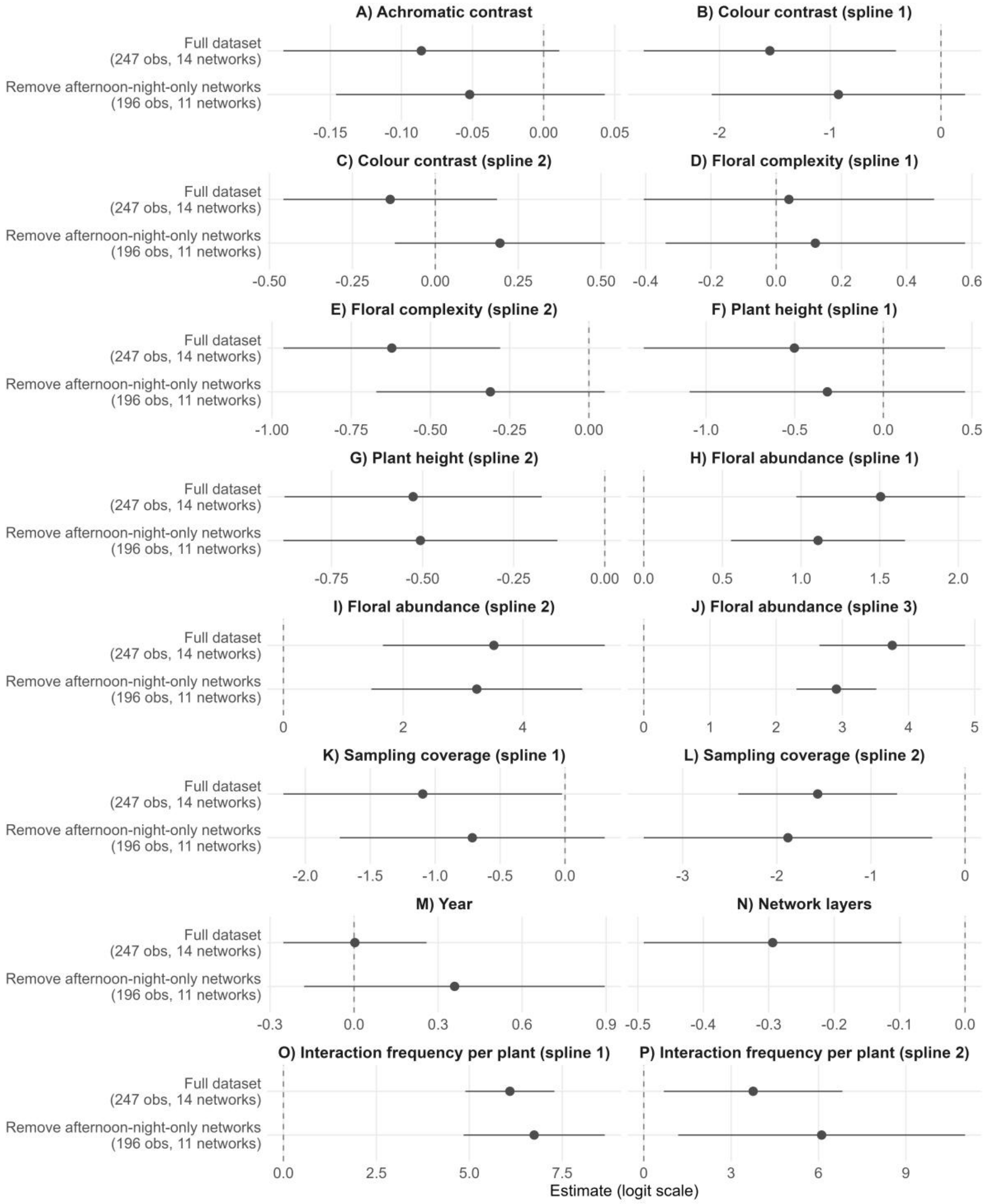

**Figure S5.** Results from the sensitivity test on the association between floral traits and versatility. Rows from top to bottom show the estimated fixed effects for the model with the full set of networks and after removing networks sampled during afternoon and night only. Estimates are indicated by the dots and lines depict 95% confidence intervals. A) Achromatic contrast, B-C) colour contrast, D-E) floral complexity, F-G) plant height, H-J) Floral abundance, K-L) Sampling coverage, M) year, N) number of network layers, O-P) interaction frequency per plant.

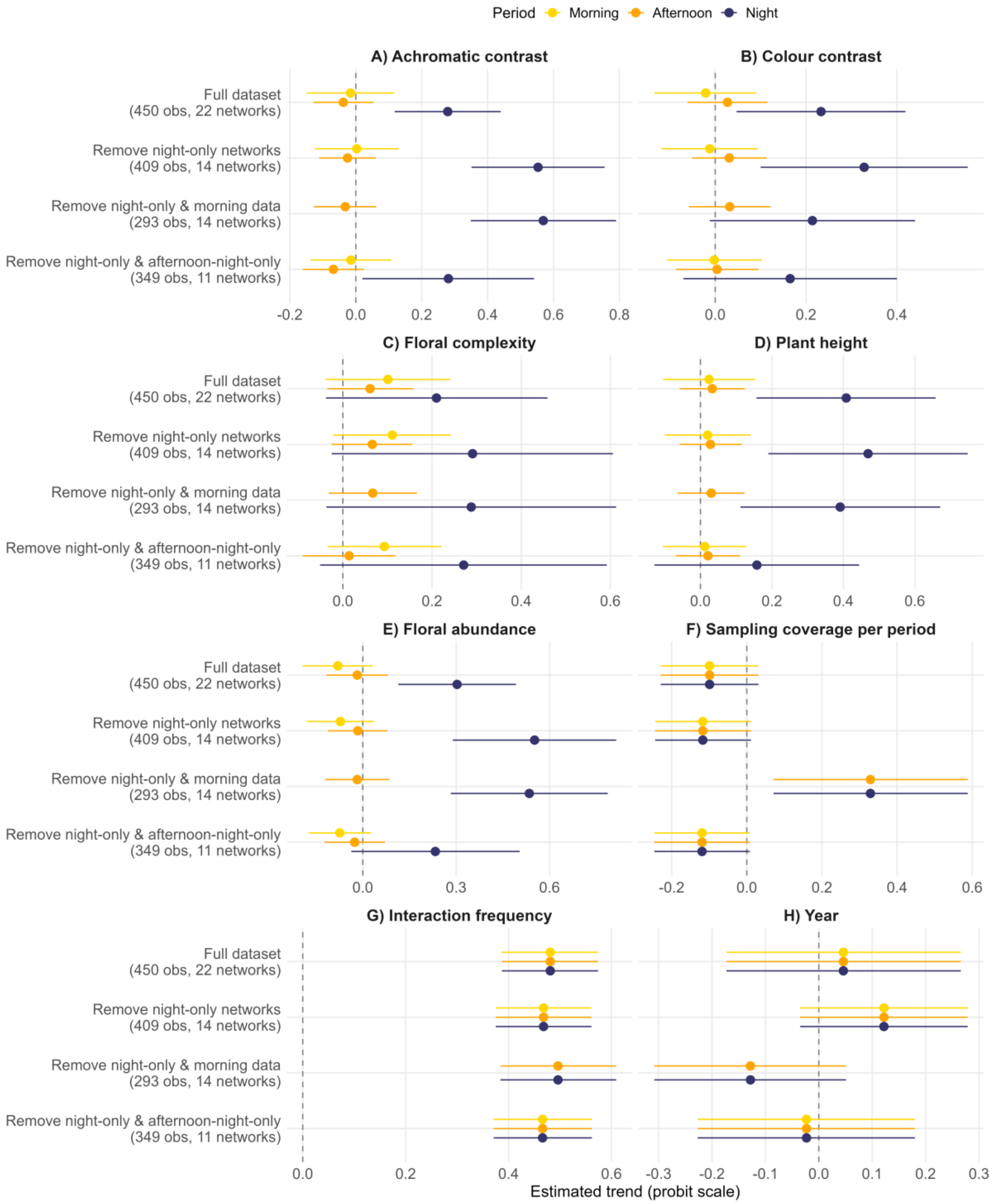

**Figure S6.** Results from the sensitivity test on the association between floral traits and plant centrality. Rows from top to bottom show the estimated trends for the model with the full set of networks and each reduced subset. Estimated trends are indicated by the dots and lines depict 95% confidence intervals. A) Achromatic contrast, B) colour contrast, C) floral complexity, D) plant height, E) floral abundance, F) sampling coverage per period, G) interaction frequency per plant and period, H) Year.

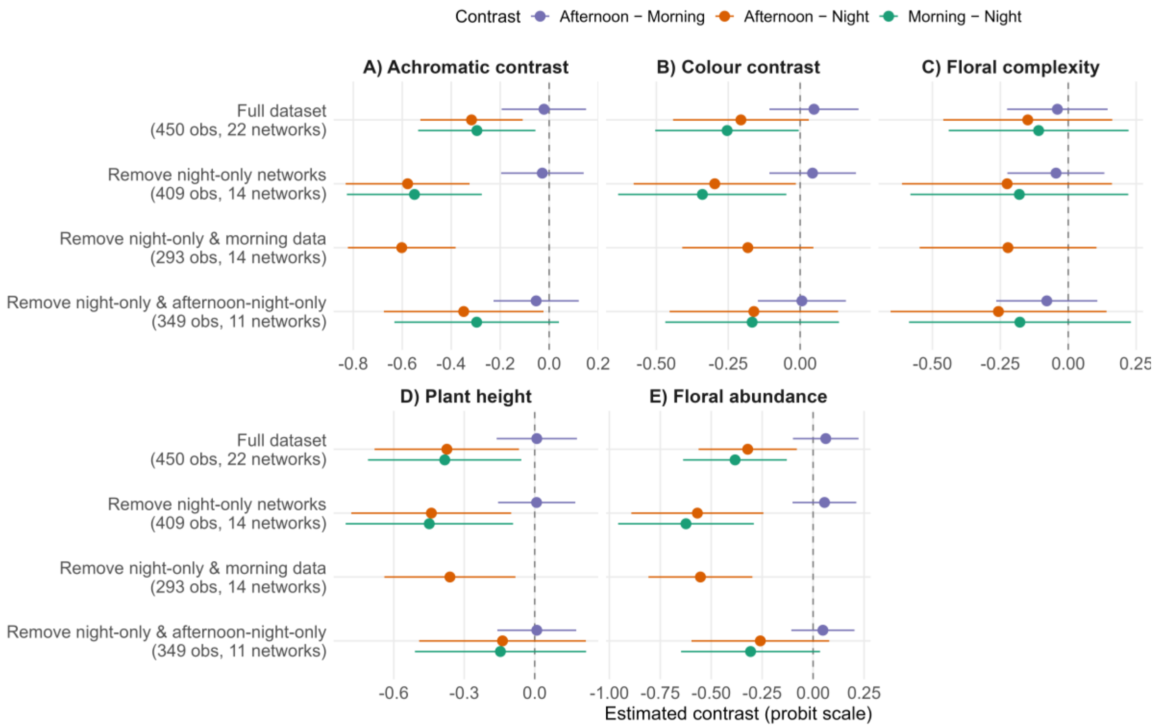

**Figure S7.** Results from the sensitivity test on pairwise contrasts between diel periods for the association between floral traits and plant centrality. Rows from top to bottom show the estimated contrasts for the model with the full set of networks and each reduced subset. Estimated contrasts are indicated by the dots and lines depict 95% confidence intervals. A) Achromatic contrast, B) colour contrast, C) floral complexity, D) plant height, E) floral abundance.

### Tables

**Table S1.** Details on the sampled sites.

| Site | Latitude | Longitude | Sampled years |
|---|---|---|---|
| A | 46.64352 | 7.5676 | 2014, 2015, 2016 |
| B | 46.66322 | 7.61897 | 2014 |
| C | 46.66005 | 7.62104 | 2014 |
| D | 46.65197 | 7.57425 | 2014 |
| E | 46.635846 | 7.56436 | 2014 |
| F | 46.646845 | 7.572463 | 2014 |
| G | 46.65611 | 7.59194 | 2014 |
| H | 46.63593 | 7.56333 | 2014 |
| I | 46.66831 | 7.61394 | 2015, 2016, 2017 |
| J | 46.630623 | 7.560865 | 2015 |
| K | 46.62537 | 7.56152 | 2015, 2016, 2017 |
| L | 46.622273 | 7.559966 | 2015 |
| M | 46.60872 | 7.521 | 2016 |
| R | 46.79235 | 7.40605 | 2016 |
| S | 46.77987 | 7.47612 | 2016, 2017 |

**Table S2.** Details on the analysed networks. Some networks were sampled only during the night, while others only during the afternoon and night, which is why their number of periods correspond to 1 and 2, respectively. Sampling coverage of interactions is shown for the full networks. Note that for network only sampled at night, this corresponds to the sampling coverage of interactions for the specific diel period.

| Number | Year | Site | Periods | Interaction frequency | Plants | Pollinators | Sampling coverage (visitors) | Sampling coverage (plants) | Sampling coverage (interactions) |
|---|---|---|---|---|---|---|---|---|---|
| 1 | 2014 | A | 3 | 316 | 27 | 97 | 0.849 | 1.000 | 0.627 |
| 2 | | B | 3 | 527 | 31 | 158 | 0.854 | 1.000 | 0.589 |
| 3 | | C | 3 | 461 | 29 | 126 | 0.855 | 1.000 | 0.623 |
| 4 | | D | 3 | 295 | 32 | 105 | 0.804 | 1.000 | 0.499 |
| 5 | | E | 3 | 251 | 23 | 114 | 0.726 | 1.000 | 0.439 |
| 6 | | F | 3 | 220 | 18 | 80 | 0.810 | 1.000 | 0.574 |
| 7 | | G | 3 | 195 | 20 | 90 | 0.688 | 1.000 | 0.483 |
| 8 | | H | 3 | 187 | 25 | 64 | 0.787 | 1.000 | 0.573 |
| 9 | 2015 | A | 1 | 112 | 7 | 31 | 0.867 | 1.000 | 0.832 |
| 10 | | I | 1 | 90 | 6 | 30 | 0.789 | 0.999 | 0.680 |
| 11 | | J | 1 | 62 | 6 | 26 | 0.777 | 1.000 | 0.614 |
| 12 | | K | 1 | 76 | 5 | 31 | 0.779 | 0.999 | 0.739 |
| 13 | | L | 1 | 20 | 4 | 10 | 0.710 | 1.000 | 0.505 |
| 14 | 2016 | A | 2 | 265 | 21 | 73 | 0.876 | 1.000 | 0.770 |
| 15 | | I | 3 | 215 | 22 | 63 | 0.824 | 0.999 | 0.680 |
| 16 | | K | 3 | 170 | 9 | 58 | 0.806 | 1.000 | 0.683 |
| 17 | | M | 3 | 499 | 25 | 112 | 0.892 | 1.000 | 0.724 |
| 18 | | R | 2 | 183 | 30 | 45 | 0.853 | 1.000 | 0.629 |
| 19 | | S | 2 | 268 | 13 | 48 | 0.903 | 1.000 | 0.806 |
| 20 | 2017 | I | 1 | 242 | 6 | 36 | 0.947 | 1.000 | 0.897 |
| 21 | | K | 1 | 97 | 6 | 22 | 0.908 | 0.990 | 0.877 |
| 22 | | S | 1 | 102 | 6 | 16 | 0.961 | 0.997 | 0.844 |

**Table S3.** Insect mouthparts. Information on mouthpart morphology was obtained from (11, 12) for most insect families and genera. This information was complemented with data from (13) for coleopterans, (14) for hemipterans, (15) for hymenopterans, and (16) for mecopterans. “Diet” indicates the insect diet based on their mouthparts (1 if they consume, 0 otherwise). N: nectar, P: pollen, T: floral tissue, S: sap, or O: other (e.g., visitors were likely looking for a prey instead of visiting the flower for typical rewards). “Type” indicates whether the taxa was considered in this study as a pollinator (Pol) or not (No).

| **Order** | **Sub order** | **Super family** | **Family** | **Sub family** | **Tribe** | **Genus** | **Diet** | | | | | **Type** |
|---|---|---|---|---|---|---|---|---|---|---|---|---|
| | | | | | | | **N** | **P** | **T** | **S** | **O** | |
| Blattodea | | | Ectobiidae | | | | 1 | 1 | 1 | 0 | 0 | No |
| Coleoptera | | | Brentidae | | | | 0 | 0 | 1 | 0 | 0 | No |
| Coleoptera | | | Buprestidae | | | | 0 | 1 | 0 | 0 | 0 | Pol |
| Coleoptera | | | Byturidae | | | | 0 | 1 | 1 | 0 | 0 | No |
| Coleoptera | | | Cantharidae | | | | 0 | 1 | 1 | 0 | 0 | No |
| Coleoptera | | | Carabidae | | | | 0 | 0 | 0 | 0 | 1 | No |
| Coleoptera | | | Cerambycidae | | | | 1 | 1 | 1 | 0 | 0 | No |
| Coleoptera | | | Chrysomelidae | | | | 0 | 1 | 1 | 0 | 0 | No |
| Coleoptera | | | Cleridae | | | | 0 | 1 | 1 | 0 | 0 | No |
| Coleoptera | | | Coccinellidae | | | | 0 | 0 | 0 | 0 | 1 | No |
| Coleoptera | | | Curculionidae | | | | 0 | 1 | 1 | 0 | 0 | No |
| Coleoptera | | | Dermestidae | | | | 0 | 1 | 0 | 0 | 0 | Pol |
| Coleoptera | | | Elateridae | | | | 0 | 1 | 1 | 0 | 0 | No |
| Coleoptera | | | Lagriidae | | | | 0 | 1 | 0 | 0 | 0 | Pol |
| Coleoptera | | | Melyridae | Dasytinae | | | 0 | 0 | 1 | 0 | 0 | No |
| Coleoptera | | | Melyridae | Malachiinae | | | 0 | 1 | 0 | 0 | 0 | Pol |
| Coleoptera | | | Melyridae | | | | 0 | 0 | 1 | 0 | 0 | No |
| Coleoptera | | | Mordellidae | | | | 0 | 1 | 1 | 0 | 0 | No |
| Coleoptera | | | Nitidulidae | | | | 0 | 1 | 1 | 0 | 0 | No |
| Coleoptera | | | Oedemeridae | | | | 1 | 1 | 1 | 0 | 0 | No |
| Coleoptera | | | Pyrochroidae | | | | 0 | 1 | 0 | 0 | 0 | Pol |
| Coleoptera | | | Scarabaeidae | Rutelinae | Anomalini | | 0 | 1 | 1 | 0 | 0 | No |
| Coleoptera | | | Scarabaeidae | Rutelinae | Hopliini | | 1 | 1 | 1 | 0 | 0 | No |
| Coleoptera | | | Scarabaeidae | | | Cetonia | 0 | 1 | 0 | 0 | 0 | Pol |
| Coleoptera | | | Scarabaeidae | | | Trichius | 0 | 1 | 0 | 0 | 0 | Pol |
| Coleoptera | | | Scarabaeidae | Cetoniinae | | | 1 | 1 | 0 | 0 | 0 | Pol |
| Coleoptera | | | Scarabaeidae | | | | 1 | 1 | 1 | 0 | 0 | No |
| Coleoptera | | | Scraptiidae | | | | 0 | 1 | 1 | 0 | 0 | No |
| Coleoptera | | | Staphylinidae | | | | 0 | 1 | 1 | 0 | 0 | No |
| Coleoptera | | | Tenebrionidae | | | | 0 | 1 | 1 | 0 | 0 | No |
| Dermaptera | | | Forficulidae | | | | 0 | 0 | 1 | 0 | 0 | No |
| Diptera | Nematocera | | NA | | | | 1 | 1 | 0 | 0 | 0 | Pol |
| Diptera | | | Anisopodidae | | | | 1 | 0 | 0 | 0 | 0 | Pol |
| Diptera | | | Anthomyiidae | | | | 1 | 1 | 0 | 0 | 0 | Pol |
| Diptera | | | Anthomyzidae | | | | 0 | 0 | 0 | 0 | 1 | No |
| Diptera | | | Asilidae | | | | 1 | 0 | 0 | 0 | 0 | Pol |

| | | | | | | | | |
|---|---|---|---|---|---|---|---|---|
| Diptera | Bibionidae | | 1 | 1 | 0 | 0 | 0 | Pol |
| Diptera | Bombyliidae | | 1 | 1 | 0 | 0 | 1 | Pol |
| Diptera | Calliphoridae | | 1 | 1 | 0 | 0 | 0 | Pol |
| Diptera | Ceratopogonidae | | 1 | 1 | 0 | 0 | 0 | Pol |
| Diptera | Chironomidae | | 1 | 0 | 0 | 0 | 0 | Pol |
| Diptera | Chloropidae | | 1 | 0 | 0 | 0 | 0 | Pol |
| Diptera | Conopidae | | 1 | 0 | 0 | 0 | 0 | Pol |
| Diptera | Culicidae | | 1 | 0 | 0 | 0 | 0 | Pol |
| Diptera | Dolichopodidae | | 1 | 0 | 0 | 0 | 1 | Pol |
| Diptera | Drosophilidae | | 1 | 1 | 0 | 0 | 0 | Pol |
| Diptera | Dryomyzidae | | 0 | 0 | 0 | 0 | 1 | No |
| Diptera | Empididae | | 1 | 1 | 0 | 0 | 1 | Pol |
| Diptera | Fanniidae | | 0 | 0 | 0 | 0 | 1 | No |
| Diptera | Hybotidae | | 0 | 0 | 0 | 0 | 1 | No |
| Diptera | Keroplatidae | | 0 | 0 | 0 | 0 | 1 | No |
| Diptera | Lauxaniidae | | 0 | 0 | 0 | 0 | 1 | No |
| Diptera | Limoniidae | | 1 | 0 | 0 | 0 | 0 | Pol |
| Diptera | Muscidae | | 1 | 1 | 0 | 0 | 0 | Pol |
| Diptera | Opomyzidae | | 0 | 0 | 0 | 0 | 1 | No |
| Diptera | Phoridae | | 1 | 0 | 0 | 0 | 1 | Pol |
| Diptera | Polleniidae | | 1 | 1 | 0 | 0 | 0 | Pol |
| Diptera | Rhagionidae | | 0 | 0 | 0 | 0 | 1 | No |
| Diptera | Rhiniidae | | 0 | 0 | 0 | 0 | 1 | No |
| Diptera | Sarcophagidae | | 1 | 1 | 0 | 0 | 0 | Pol |
| Diptera | Scathophagidae | | 1 | 1 | 0 | 0 | 1 | Pol |
| Diptera | Scatopsidae | | 0 | 1 | 0 | 0 | 0 | Pol |
| Diptera | Sciaridae | | 1 | 0 | 0 | 0 | 0 | Pol |
| Diptera | Sciomyzidae | | 0 | 0 | 0 | 0 | 1 | No |
| Diptera | Sepsidae | | 1 | 0 | 0 | 0 | 0 | Pol |
| Diptera | Sphaeroceridae | | 0 | 0 | 0 | 0 | 1 | No |
| Diptera | Stratiomyidae | | 0 | 0 | 0 | 0 | 1 | No |
| Diptera | Syrphidae | | 1 | 1 | 0 | 0 | 0 | Pol |
| Diptera | Tabanidae | | 1 | 0 | 0 | 0 | 1 | Pol |
| Diptera | Tachinidae | | 1 | 1 | 0 | 0 | 0 | Pol |
| Diptera | Tephritidae | | 1 | 0 | 0 | 0 | 1 | Pol |
| Diptera | Therevidae | | 0 | 0 | 0 | 0 | 1 | No |
| Diptera | Tipulidae | | 1 | 0 | 0 | 0 | 1 | Pol |
| Hemiptera | Alydidae | | 0 | 0 | 0 | 1 | 0 | No |
| Hemiptera | Anthocoridae | Anthocoris | 0 | 0 | 0 | 0 | 1 | No |
| Hemiptera | Anthocoridae | Orius | 0 | 0 | 0 | 0 | 1 | No |
| Hemiptera | Aphididae | | 0 | 0 | 0 | 1 | 0 | No |
| Hemiptera | Aphrophoridae | | 0 | 0 | 0 | 1 | 0 | No |
| Hemiptera | Caliscelidae | | 0 | 0 | 0 | 1 | 0 | No |
| Hemiptera | Cercopidae | | 0 | 0 | 0 | 1 | 0 | No |

| | | | | | | | | | | |
|---|---|---|---|---|---|---|---|---|---|---|
| Hemiptera | | Cicadellidae | | | 0 | 0 | 0 | 1 | 0 | No |
| Hemiptera | | Coreidae | | | 0 | 0 | 0 | 1 | 0 | No |
| Hemiptera | | Lygaeidae | | | 0 | 0 | 0 | 1 | 0 | No |
| Hemiptera | | Miridae | | Adelphocoris | 0 | 0 | 0 | 1 | 1 | No |
| Hemiptera | | Miridae | | Calocoris | 0 | 1 | 0 | 1 | 0 | No |
| Hemiptera | | Miridae | | Closterotomus | 0 | 0 | 0 | 1 | 0 | No |
| Hemiptera | | Miridae | | Deraeocoris | 0 | 0 | 0 | 1 | 1 | No |
| Hemiptera | | Miridae | | Dicyphus | 0 | 0 | 0 | 0 | 1 | No |
| Hemiptera | | Miridae | | Globiceps | 0 | 0 | 0 | 1 | 1 | No |
| Hemiptera | | Miridae | | Grypocoris | 0 | 0 | 0 | 1 | 1 | No |
| Hemiptera | | Miridae | | Liocoris | 0 | 0 | 0 | 1 | 0 | No |
| Hemiptera | | Miridae | | Lygocoris | 0 | 0 | 0 | 1 | 1 | No |
| Hemiptera | | Miridae | | Lygus | 1 | 0 | 0 | 1 | 1 | No |
| Hemiptera | | Miridae | | Orthops | 0 | 0 | 0 | 1 | 0 | No |
| Hemiptera | | Miridae | | Phytocoris | 0 | 0 | 0 | 1 | 1 | No |
| Hemiptera | | Miridae | | Plagiognathus | 1 | 0 | 0 | 1 | 1 | No |
| Hemiptera | | Miridae | | Psallus | 0 | 0 | 0 | 1 | 1 | No |
| Hemiptera | | Miridae | | Stenodema | 0 | 0 | 0 | 1 | 0 | No |
| Hemiptera | | Miridae | | Stenodema | 0 | 0 | 0 | 1 | 0 | No |
| Hemiptera | | Miridae | | Stenotus | 0 | 0 | 0 | 1 | 0 | No |
| Hemiptera | | Miridae | Phylinae | | 0 | 0 | 0 | 1 | 1 | No |
| Hemiptera | | Miridae | | | 0 | 0 | 0 | 1 | 1 | No |
| Hemiptera | | Nabidae | | | 0 | 0 | 0 | 0 | 1 | No |
| Hemiptera | | Pentatomidae | Asopinae | | 0 | 0 | 0 | 0 | 1 | No |
| Hemiptera | | Pentatomidae | | | 0 | 0 | 0 | 1 | 0 | No |
| Hemiptera | | Pyrrhocoridae | | | 0 | 0 | 0 | 1 | 0 | No |
| Hemiptera | | Rhopalidae | | | 1 | 0 | 0 | 1 | 0 | No |
| Hemiptera | | Rhyparochromidae | | | 0 | 0 | 0 | 1 | 0 | No |
| Hemiptera | | Scutelleridae | | | 0 | 0 | 0 | 1 | 0 | No |
| Hymenoptera | Apoidea | Andrenidae | | | 1 | 0 | 0 | 0 | 0 | Pol |
| Hymenoptera | Apoidea | Apidae | | | 1 | 1 | 0 | 0 | 0 | Pol |
| Hymenoptera | Apoidea | Colletidae | | | 1 | 0 | 0 | 0 | 0 | Pol |
| Hymenoptera | Apoidea | Crabronidae | | | 0 | 0 | 0 | 0 | 1 | No |
| Hymenoptera | Apoidea | Halictidae | | | 1 | 0 | 0 | 0 | 0 | Pol |
| Hymenoptera | Apoidea | Megachilidae | | | 1 | 0 | 0 | 0 | 0 | Pol |
| Hymenoptera | Apoidea | Melittidae | | | 1 | 0 | 0 | 0 | 0 | Pol |
| Hymenoptera | Apoidea | Sphecidae | | | 1 | 0 | 0 | 0 | 0 | Pol |
| Hymenoptera | Chrysidoidea | Chrysididae | | | 1 | 0 | 0 | 0 | 0 | Pol |
| Hymenoptera | Formicoidea | Formicidae | | | 1 | 0 | 1 | 0 | 1 | No |
| Hymenoptera | Ichneumonoidea | Braconidae | | | 1 | 0 | 0 | 0 | 0 | Pol |
| Hymenoptera | Ichneumonoidea | Ichneumonidae | | | 1 | 0 | 0 | 0 | 0 | Pol |

| | | | | | | | | | |
|---|---|---|---|---|---|---|---|---|---|
| Hymenoptera | Pamphilioidea | Pamphiliidae | | 0 | 0 | 0 | 0 | 1 | No |
| Hymenoptera | Pompiloidea | Pompilidae | | 1 | 0 | 0 | 0 | 0 | Pol |
| Hymenoptera | Tenthredinoidea | Argidae | | 0 | 0 | 0 | 0 | 1 | No |
| Hymenoptera | Tenthredinoidea | Pergidae | | 1 | 0 | 0 | 0 | 0 | Pol |
| Hymenoptera | Tenthredinoidea | Tenthredinidae | | 1 | 0 | 1 | 0 | 0 | No |
| Hymenoptera | Tiphioidea | Tiphiidae | | 1 | 0 | 0 | 0 | 0 | Pol |
| Hymenoptera | Vespoidea | Vespidae | Vespinae | 1 | 0 | 0 | 0 | 0 | Pol |
| Hymenoptera | Vespoidea | Vespidae | Eumeninae | 1 | 0 | 0 | 0 | 0 | Pol |
| Hymenoptera | Vespoidea | Vespidae | Polistinae | 1 | 0 | 0 | 0 | 0 | Pol |
| Lepidoptera | | Adelidae | | 1 | 0 | 0 | 0 | 0 | Pol |
| Lepidoptera | | Arctiidae | | 1 | 0 | 0 | 0 | 0 | Pol |
| Lepidoptera | | Crambidae | | 1 | 0 | 0 | 0 | 0 | Pol |
| Lepidoptera | | Erebidae | | 1 | 0 | 0 | 0 | 0 | Pol |
| Lepidoptera | | Geometridae | | 1 | 0 | 0 | 0 | 0 | Pol |
| Lepidoptera | | Hesperiidae | | 1 | 0 | 0 | 0 | 0 | Pol |
| Lepidoptera | | Lycaenidae | | 1 | 0 | 0 | 0 | 0 | Pol |
| Lepidoptera | | NA | | 1 | 0 | 0 | 0 | 0 | Pol |
| Lepidoptera | | Noctuidae | | 1 | 0 | 0 | 0 | 0 | Pol |
| Lepidoptera | | Nolidae | | 1 | 0 | 0 | 0 | 0 | Pol |
| Lepidoptera | | Nymphalidae | | 1 | 0 | 0 | 0 | 0 | Pol |
| Lepidoptera | | Pieridae | | 1 | 0 | 0 | 0 | 0 | Pol |
| Lepidoptera | | Pterophoridae | | 1 | 0 | 0 | 0 | 0 | Pol |
| Lepidoptera | | Pyralidae | | 1 | 0 | 0 | 0 | 0 | Pol |
| Lepidoptera | | Riodinidae | | 1 | 0 | 0 | 0 | 0 | Pol |
| Lepidoptera | | Sphingidae | | 1 | 0 | 0 | 0 | 0 | Pol |
| Lepidoptera | | Tortricidae | | 1 | 0 | 0 | 0 | 0 | Pol |
| Lepidoptera | | Zygaenidae | | 1 | 0 | 0 | 0 | 0 | Pol |
| Mecoptera | | Panorpidae | | 1 | 1 | 0 | 0 | 1 | Pol |
| Neuroptera | | Chrysopidae | | 1 | 1 | 0 | 0 | 0 | Pol |
| Neuroptera | | Hemerobiidae | | 1 | 1 | 1 | 0 | 0 | No |
| Orthoptera | | Acrididae | | 0 | 0 | 1 | 0 | 0 | No |
| Orthoptera | | Tetrigidae | | 0 | 0 | 1 | 0 | 0 | No |
| Orthoptera | | Tettigoniidae | | 0 | 0 | 1 | 0 | 0 | No |

**Table S4.** Floral traits used in our analysis. Reflectance values were not available for all plant species; therefore, we were not able to estimate achromatic and chromatic contrast for all of them. FCI: Floral complexity index.

| Plant | FCI | Achromatic contrast | Chromatic contrast | Plant height (cm) |
|---|---|---|---|---|
| *Achillea millefolium* | 1.56 | 0.301445037 | 0.34231621 | 80 |
| *Aconitum lycoctonum* | 2.9 | NA | NA | 180 |
| *Aegopodium podagraria* | 1.18 | 0.245538194 | 0.2152252 | 90 |
| *Ajuga reptans* | 2.83 | 0.008910961 | 0.24654322 | 30 |
| *Alchemilla xanthochlora* | 1.33 | NA | NA | 31 |
| *Alliaria petiolata* | 1.91 | 0.335624757 | 0.12227335 | 90 |
| *Angelica sylvestris* | 1.18 | 0.326283353 | 0.21119984 | 150 |
| *Anthriscus sylvestris* | 1.55 | 0.306264633 | 0.32966405 | 150 |
| *Anthyllis vulneraria* | 2.83 | 0.246949148 | 0.15631461 | 60 |
| *Apiaceae* | NA | NA | NA | NA |
| *Aquilegia atrata* | 2.18 | NA | NA | 90 |
| *Arctium minus* | 1.56 | NA | NA | 130 |
| *Aruncus dioicus* | 2.08 | NA | NA | 200 |
| *Asteraceae* | NA | NA | NA | NA |
| *Astrantia major* | 1.56 | 0.31941776 | 0.12283076 | 90 |
| *Bellis perennis* | 1.56 | 0.335988324 | 0.16103262 | 15 |
| *Campanula* sp. | 2.53 | 0.073763998 | 0.221055 | 100 |
| *Campanula trachelium* | 2.53 | 0.073763998 | 0.221055 | 100 |
| *Cardamine impatiens* | 2.08 | NA | NA | 80 |
| *Carduus personata* | 1.56 | NA | NA | 150 |
| *Centaurea jacea* | 1.96 | 0.076705144 | 0.19186264 | 80 |
| *Centaurea montana* | 1.76 | 0.110169827 | 0.17680495 | 60 |
| *Centaurea* sp. | 1.96 | NA | NA | 70 |
| *Chaerophyllum hirsutum* | 1.18 | NA | NA | 73 |
| *Circaea lutetiana* | 1.33 | NA | NA | 70 |
| *Cirsium arvense* | 1.56 | 0.086044041 | 0.27173564 | 100 |
| *Cirsium eriophorum* | 1.56 | NA | NA | 150 |
| *Cirsium oleraceum* | 1.56 | 0.239269091 | 0.27931027 | 150 |
| *Cirsium palustre* | 1.56 | 0.049244129 | 0.29908765 | 150 |
| *Cirsium vulgare* | 1.76 | NA | NA | 150 |
| *Cornus sanguinea* | 1.93 | 0.284351434 | 0.21605014 | 200 |
| *Crepis biennis* | 1.56 | NA | NA | 100 |
| *Crepis paludosa* | 1.56 | 0.268874297 | 0.20347363 | 120 |
| *Crepis sp* | 1.56 | 0.268874297 | 0.20347363 | 60 |
| *Daucus carota* | 1.18 | 0.405672583 | 0.216878 | 100 |
| *Epilobium angustifolium* | 1.33 | 0.058152129 | 0.26073606 | 150 |
| *Epilobium hirsutum* | 2.23 | 0.01199421 | 0.2566149 | 150 |
| *Epilobium parviflorum* | 2.03 | 0.176889721 | 0.20938896 | 80 |
| *Epilobium* sp. | 2.03 | NA | NA | 124 |
| *Erigeron annuus* | 1.56 | NA | NA | 150 |
| *Eupatorium cannabinum* | 1.93 | 0.077783591 | 0.26643749 | 150 |

| | | | | |
|---|---|---|---|---|
| *Filipendula ulmaria* | 1.93 | 0.303035195 | 0.24619971 | 200 |
| *Fragaria vesca* | 1.18 | 0.321690476 | 0.17539733 | 20 |
| *Galeopsis tetrahit* | 3.03 | 0.176368476 | 0.26236976 | 100 |
| *Galium mollugo* | 1.33 | 0.246887818 | 0.23850272 | 77 |
| *Galium odoratum* | 1.96 | NA | NA | 30 |
| *Galium sylvaticum* | 1.33 | NA | NA | 80 |
| *Galium verum* | 1.33 | 0.199224186 | 0.41030804 | 70 |
| *Geranium pyrenaicum* | 1.18 | 0.176748902 | 0.19445373 | 60 |
| *Geranium robertianum* | 1.38 | 0.14295572 | 0.15369016 | 50 |
| *Geranium sylvaticum* | 1.18 | NA | NA | 60 |
| *Geum rivale* | 2.28 | 0.073558585 | 0.13819601 | 60 |
| *Geum urbanum* | 1.18 | 0.302741179 | 0.28445691 | 90 |
| *Glechoma hederacea* | 3.03 | 0.001886701 | 0.25174183 | 24 |
| *Heracleum sphondylium* | 1.18 | 0.335893917 | 0.2666312 | 150 |
| *Hieracium* sp. | 1.71 | NA | NA | NA |
| *Hippocrepis comosa* | 2.83 | NA | NA | 25 |
| *Hypericum maculatum* | 2.08 | 0.261292673 | 0.20116901 | 60 |
| *Hypericum perforatum* | 2.08 | 0.21565626 | 0.36912264 | 100 |
| *Hypochaeris radicata* | 1.56 | 0.249385976 | 0.22667307 | 60 |
| *Impatiens noli-tangere* | 2.88 | 0.260494174 | 0.12135924 | 100 |
| *Impatiens parviflora* | 2.83 | 0.314965512 | 0.07553662 | 100 |
| *Jacobaea vulgaris* | 1.56 | 0.236254815 | 0.25128395 | 100 |
| *Knautia dipsacifolia* | 1.86 | 0.146922616 | 0.29514509 | 100 |
| *Lamium album* | 3.03 | 0.181490058 | 0.26534948 | 33 |
| *Lamium galeobdolon* | 3.03 | 0.283377247 | 0.11482904 | 60 |
| *Lamium maculatum* | 3.03 | 0.081047857 | 0.15432915 | 50 |
| *Lamium purpureum* | 3.03 | 0.098219051 | 0.17649875 | 25 |
| *Lapsana communis* | | 0.244244368 | 0.289159766 | 120 |
| *Lathyrus pratensis* | 3.18 | 0.199902234 | 0.38482261 | 90 |
| *Leontodon sp* | 1.76 | 0.146584559 | 0.13077884 | 60 |
| *Leucanthemum vulgare* | 1.56 | 0.364018829 | 0.21279114 | 46 |
| *Lonicera ligustrina* | 2.23 | NA | NA | NA |
| *Lotus corniculatus* | 2.83 | 0.186600804 | 0.44155091 | 30 |
| *Malva moschata* | 1.33 | 0.138607788 | 0.15762325 | 60 |
| *Matricaria discoidea* | | 0.114504738 | 0.063905493 | 19 |
| *Medicago lupulina* | 2.63 | 0.215371912 | 0.24061341 | 30 |
| *Melilotus albus* | 2.98 | NA | NA | 150 |
| *Mentha longifolia* | 2.55 | 0.169282221 | 0.26348105 | 100 |
| *Myosotis scorpioides* | 1.96 | NA | NA | 50 |
| *Myosotis* sp. | 1.96 | NA | NA | 50 |
| *Oenothera biennis* | 1.83 | 0.29566205 | 0.15717911 | 150 |
| *Oxalis acetosella* | 1.18 | NA | NA | 15 |
| *Persicaria bistorta* | 2.38 | 0.249777579 | 0.19179453 | 80 |
| *Persicaria maculosa* | 1.43 | NA | NA | 46 |

| | | | | |
|---|---|---|---|---|
| *Phyteuma spicatum* | 2.61 | NA | NA | 70 |
| *Plantago lanceolata* | 1.96 | NA | NA | 40 |
| *Potentilla erecta* | 1.18 | 0.263320119 | 0.41192257 | 60 |
| *Potentilla reptans* | 1.18 | 0.249333045 | 0.19499402 | 100 |
| *Prunella vulgaris* | 3.03 | 0.039326095 | 0.17384619 | 20 |
| *Ranunculus* sp. | 1.18 | 0.27015517 | 0.31842336 | 100 |
| *Rorippa sylvestris* | 1.33 | NA | NA | 60 |
| *Rubus caesius* | 2.08 | 0.296552575 | 0.24402925 | 68 |
| *Rubus fruticosus* | 2.08 | NA | NA | 200 |
| *Rubus idaeus* | 2.08 | NA | NA | 150 |
| *Rubus* sp. | 2.08 | NA | NA | 172.6667 |
| *Salvia glutinosa* | 3.03 | NA | NA | 100 |
| *Salvia pratensis* | 3.03 | 0.044940575 | 0.24071778 | 60 |
| *Scorzoneroides autumnalis* | 1.56 | NA | NA | 50 |
| *Silene dioica* | 2.36 | 0.087466912 | 0.20341869 | 90 |
| *Silene vulgaris* | 2.36 | 0.309512442 | 0.27198662 | 50 |
| *Solidago canadensis* | 1.71 | 0.208141451 | 0.4088681 | 200 |
| *Spiraea japonica* | 1.93 | NA | NA | 192 |
| *Stachys alpina* | 3.03 | NA | NA | 59 |
| *Stachys sylvatica* | 3.03 | 0.027478164 | 0.12686174 | 100 |
| *Stellaria media* | 1.18 | NA | NA | 40 |
| *Symphoricarpos albus* | 2.33 | 0.140074215 | 0.12416019 | 200 |
| *Symphytum officinale* | 2.61 | 0.338219039 | 0.16089022 | 120 |
| *Syringa vulgaris* | 2.73 | 0.26857207 | 0.08679226 | 200 |
| *Taraxacum officinale* | 1.56 | 0.225097821 | 0.36198863 | 30 |
| *Taraxacum palustre* | 1.56 | 0.225097821 | 0.36198863 | 30 |
| *Thymus* sp. | 2.68 | 0.151615575 | 0.1471031 | 25 |
| *Trifolium pratense* | 2.98 | 0.005253382 | 0.22927785 | 45 |
| *Trifolium repens* | 2.83 | 0.324359693 | 0.18720053 | 40 |
| *Urtica dioica* | 1.33 | NA | NA | 100 |
| *Valeriana officinalis* | 1.78 | 0.27520498 | 0.12558247 | 160 |
| *Verbena officinalis* | 2.5 | NA | NA | 70 |
| *Veronica chamaedrys* | 2.33 | 0.040081785 | 0.23466229 | 30 |
| *Veronica* sp. | 2.33 | 0.040081785 | 0.23466229 | 30 |
| *Viburnum opulus* | 1.18 | 0.335211356 | 0.15260353 | 200 |
| *Vicia cracca* | 3.18 | 0.057793293 | 0.27349325 | 120 |
| *Vicia lutea* | 3.03 | NA | NA | 50 |
| *Vicia sepium* | 3.18 | 0.158503658 | 0.08706348 | 90 |

**Table S5.** Floral traits used to calculate the Floral Complexity index (FCI).

| Plant species | Floral shape | Floral depth | Floral symmetry | Corolla segmentation | Functional reproductive unit | Corolla tube length (mm) | Corolla tube length source |
|---|---|---|---|---|---|---|---|
| *Achillea millefolium* | head | low | radial | choripetaly | flat or spherical | 1.37 | (17) via TRY (18) |
| *Aconitum lycoctonum* | bell | high | bilateral | semichoripetaly | cylindrical | 35.5 | (19) |
| *Aegopodium podagraria* | disk | low | radial | choripetaly | flat or spherical | 0 | (20) |
| *Ajuga reptans* | gullet | medium | bilateral | semichoripetaly | cylindrical | 8.46 | (17) via TRY (18) |
| *Alchemilla xanthochlora* | disk | low | radial | choripetaly | cylindrical | 1.45 | (17) via TRY (18) |
| *Alliaria petiolata* | disk-tube | medium | radial | choripetaly | flat or spherical | 6.5 | (21) |
| *Angelica sylvestris* | disk | low | radial | choripetaly | flat or spherical | NA | |
| *Anthriscus sylvestris* | disk | low | bilateral | choripetaly | flat or spherical | 0 | (22) via FLoRes (23) |
| *Anthyllis vulneraria* | flag | medium | bilateral | semichoripetaly | flat or spherical | 4 | (24) |
| *Aquilegia atrata* | bell | medium | radial | semichoripetaly | single | NA | |
| *Arctium minus* | head | low | radial | choripetaly | flat or spherical | 3.9 | (25) via a FloRes (23) |
| *Aruncus dioicus* | brush | low | radial | choripetaly | cylindrical | NA | |
| *Astrantia major* | head | low | radial | choripetaly | flat or spherical | NA | |
| *Bellis perennis* | head | low | radial | choripetaly | flat or spherical | 1.27 | (17) via TRY (18) |
| *Campanula sp* | bell | high | radial | semichoripetaly | cylindrical | NA | |
| *Campanula trachelium* | bell | high | radial | semichoripetaly | cylindrical | 32.5 | (26) via InfoFlora (https://www.infoflora.ch/) |
| *Cardamine impatiens* | brush | low | radial | choripetaly | cylindrical | NA | |
| *Carduus personata* | head | low | radial | choripetaly | flat or spherical | NA | |
| *Centaurea jacea* | head | high | radial | choripetaly | flat or spherical | 12.06 | (17) via TRY (18) |
| *Centaurea montana* | head | medium | radial | choripetaly | flat or spherical | 7.2 | (27) |
| *Centaurea sp* | head | high | radial | choripetaly | flat or spherical | NA | |
| *Chaerophyllum hirsutum* | disk | low | radial | choripetaly | flat or spherical | NA | |
| *Circaea lutetiana* | disk | low | radial | choripetaly | cylindrical | NA | |
| *Cirsium arvense* | head | low | radial | choripetaly | flat or spherical | 0.96 | (28) |
| *Cirsium eriophorum* | head | low | radial | choripetaly | flat or spherical | NA | |

| | | | | | | | |
|---|---|---|---|---|---|---|---|
| *Cirsium oleraceum* | head | low | radial | choripetaly | flat or spherical | NA | |
| *Cirsium palustre* | head | low | radial | choripetaly | flat or spherical | 3 | (25) via a FloRes (23) |
| *Cirsium vulgare* | head | medium | radial | choripetaly | flat or spherical | 6.2 | (25) via a FloRes (23) |
| *Cornus sanguinea* | brush | low | radial | choripetaly | flat or spherical | NA | |
| *Crepis biennis* | head | low | radial | choripetaly | flat or spherical | NA | |
| *Crepis paludosa* | head | low | radial | choripetaly | flat or spherical | NA | |
| *Crepis sp* | head | low | radial | choripetaly | flat or spherical | NA | |
| *Daucus carota* | disk | low | radial | choripetaly | flat or spherical | 0 | (20) |
| *Epilobium angustifolium* | disk | low | radial | choripetaly | cylindrical | 0 | (29) |
| *Epilobium hirsutum* | funnel | high | radial | choripetaly | cylindrical | NA | |
| *Epilobium parviflorum* | funnel | medium | radial | choripetaly | cylindrical | 8 | FloraWeb (www.floraweb.de) |
| *Epilobium sp* | funnel | medium | radial | choripetaly | cylindrical | NA | |
| *Erigeron annuus* | head | low | radial | choripetaly | flat or spherical | NA | |
| *Eupatorium cannabinum* | brush | low | radial | choripetaly | flat or spherical | 2.71 | (30) |
| *Filipendula ulmaria* | brush | low | radial | choripetaly | flat or spherical | NA | |
| *Fragaria vesca* | disk | low | radial | choripetaly | single | 0 | (17) via TRY (18) |
| *Galeopsis tetrahit* | gullet | high | bilateral | semichoripetaly | cylindrical | 12.51 | (17) via TRY (18) |
| *Galium mollugo* | disk | low | radial | choripetaly | cylindrical | 0.86 | (17) via TRY (18) |
| *Galium odoratum* | disk-tube | low | radial | semichoripetaly | cylindrical | 1 | (26) via InfoFlora (https://www.infoflora.ch/) |
| *Galium sylvaticum* | disk | low | radial | choripetaly | cylindrical | NA | |
| *Galium verum* | disk | low | radial | choripetaly | cylindrical | NA | |
| *Geranium pyrenaicum* | disk | low | radial | choripetaly | single | NA | |
| *Geranium robertianum* | disk | medium | radial | choripetaly | single | 6.12 | (30) |
| *Geranium sylvaticum* | disk | low | radial | choripetaly | single | 0 | (17) via TRY (18) |
| *Geum rivale* | bell | high | radial | choripetaly | single | 12.5 | FloraWeb (www.floraweb.de) |
| *Geum urbanum* | disk | low | radial | choripetaly | single | NA | |
| *Glechoma hederacea* | gullet | high | bilateral | semichoripetaly | cylindrical | 15 | FloraWeb (www.floraweb.de) |

| | | | | | | | |
|---|---|---|---|---|---|---|---|
| *Heracleum sphondylium* | disk | low | radial | choripetaly | flat or spherical | 0 | (20) |
| *Hieracium sp* | head | low | radial | choripetaly | cylindrical | NA | |
| *Hippocrepis comosa* | flag | medium | bilateral | semichoripetaly | flat or spherical | NA | |
| *Hypericum maculatum* | brush | low | radial | choripetaly | cylindrical | 0 | (17) via TRY (18) |
| *Hypericum perforatum* | brush | low | radial | choripetaly | cylindrical | NA | |
| *Hypochaeris radicata* | head | low | radial | choripetaly | flat or spherical | NA | |
| *Impatiens noli-tangere* | gullet | high | bilateral | semichoripetaly | single | 35 | (26) via InfoFlora (https://www.infoflora.ch/) |
| *Impatiens parviflora* | gullet | medium | bilateral | semichoripetaly | cylindrical | 10 | (26) via InfoFlora (https://www.infoflora.ch/) |
| *Jacobaea vulgaris* | head | low | radial | choripetaly | flat or spherical | 3.72 | (30) |
| *Knautia dipsacifolia* | head | medium | radial | semichoripetaly | flat or spherical | NA | |
| *Lamium album* | gullet | high | bilateral | semichoripetaly | cylindrical | 22.5 | (26) via InfoFlora (https://www.infoflora.ch/) |
| *Lamium galeobdolon* | gullet | high | bilateral | semichoripetaly | cylindrical | 18.5 | (26) via InfoFlora (https://www.infoflora.ch/) |
| *Lamium maculatum* | gullet | high | bilateral | semichoripetaly | cylindrical | 25 | (26) via InfoFlora (https://www.infoflora.ch/) |
| *Lamium purpureum* | gullet | high | bilateral | semichoripetaly | cylindrical | 10 | (26) via InfoFlora (https://www.infoflora.ch/) |
| *Lapsana communis* | head | low | radial | choripetaly | single | NA | |
| *Lathyrus pratensis* | flag | high | bilateral | semichoripetaly | cylindrical | 12 | (26) via InfoFlora (https://www.infoflora.ch/) |
| *Leontodon sp* | head | medium | radial | choripetaly | flat or spherical | 5.35 | (17) via TRY (18) (from *L. hispidus*) |
| *Leucanthemum vulgare* | head | low | radial | choripetaly | flat or spherical | 2.13 | (17) via TRY (18) |
| *Lonicera ligustrina* | funnel | high | radial | semichoripetaly | cylindrical | 12.38 | (31) (from *Lonicera* spp.) |
| *Lotus corniculatus* | flag | medium | bilateral | semichoripetaly | flat or spherical | 7.49 | (17) via TRY (18) |
| *Malva moschata* | disk | low | radial | choripetaly | cylindrical | NA | |
| *Matricaria discoidea* | disk | low | radial | choripetaly | single | NA | |
| *Medicago lupulina* | flag | low | bilateral | semichoripetaly | flat or spherical | 2.75 | (26) via InfoFlora (https://www.infoflora.ch/) |
| *Melilotus albus* | flag | medium | bilateral | semichoripetaly | cylindrical | 4.5 | (26) via InfoFlora (https://www.infoflora.ch/) |

| | | | | | | | |
|---|---|---|---|---|---|---|---|
| *Mentha longifolia* | brush | low | bilateral | semichoripetaly | cylindrical | 3.5 | (26) via InfoFlora (https://www.infoflora.ch/) |
| *Myosotis scorpioides* | disk-tube | low | radial | semichoripetaly | cylindrical | 1 | FloraWeb (www.floraweb.de) |
| *Myosotis sp* | disk-tube | low | radial | semichoripetaly | cylindrical | NA | |
| *Oenothera biennis* | funnel | low | radial | choripetaly | cylindrical | NA | |
| *Oxalis acetosella* | disk | low | radial | choripetaly | single | NA | |
| *Persicaria bistorta* | brush | medium | radial | semichoripetaly | cylindrical | 4.5 | (26) via InfoFlora (https://www.infoflora.ch/) |
| *Persicaria maculosa* | disk | low | radial | semichoripetaly | cylindrical | NA | |
| *Phyteuma spicatum* | tube | high | radial | sympetaly | cylindrical | 12.5 | (26) via InfoFlora (https://www.infoflora.ch/) |
| *Plantago lanceolata* | disk-tube | low | radial | semichoripetaly | cylindrical | 3 | (26) via InfoFlora (https://www.infoflora.ch/) |
| *Potentilla erecta* | disk | low | radial | choripetaly | single | 0 | (17) via TRY (18) |
| *Potentilla reptans* | disk | low | radial | choripetaly | single | NA | |
| *Prunella vulgaris* | gullet | high | bilateral | semichoripetaly | cylindrical | 12.5 | (26) via InfoFlora (https://www.infoflora.ch/) |
| *Ranunculus sp* | disk | low | radial | choripetaly | single | 0 | (17) via TRY (18) (from *R. acris*) |
| *Rorippa sylvestris* | disk | low | radial | choripetaly | cylindrical | NA | |
| *Rubus caesius* | brush | low | radial | choripetaly | cylindrical | NA | |
| *Rubus fruticosus* | brush | low | radial | choripetaly | cylindrical | 0 | (30) |
| *Rubus idaeus* | brush | low | radial | choripetaly | cylindrical | NA | |
| *Rubus sp* | brush | low | radial | choripetaly | cylindrical | NA | |
| *Salvia glutinosa* | gullet | high | bilateral | semichoripetaly | cylindrical | 37.5 | (26) via InfoFlora (https://www.infoflora.ch/) |
| *Salvia pratensis* | gullet | high | bilateral | semichoripetaly | cylindrical | 22.5 | (26) via InfoFlora (https://www.infoflora.ch/) |
| *Scorzoneroides autumnalis* | head | low | radial | choripetaly | flat or spherical | NA | |
| *Silene dioica* | disk-tube | high | radial | semichoripetaly | cylindrical | 13.3 | (17) via TRY (18) |
| *Silene vulgaris* | disk-tube | high | radial | semichoripetaly | cylindrical | 13.83 | (17) via TRY (18) |
| *Solidago canadensis* | head | low | radial | choripetaly | cylindrical | 2 | (29) |
| *Spiraea japonica* | brush | low | radial | choripetaly | flat or spherical | NA | |

| | | | | | | | |
|---|---|---|---|---|---|---|---|
| *Stachys alpina* | gullet | high | bilateral | semichoripetaly | cylindrical | 16.5 | (26) via InfoFlora (https://www.infoflora.ch/) |
| *Stachys sylvatica* | gullet | high | bilateral | semichoripetaly | cylindrical | 13.5 | (26) via InfoFlora (https://www.infoflora.ch/) |
| *Stellaria media* | disk | low | radial | choripetaly | single | NA | |
| *Symphoricarpos albus* | bell | medium | radial | semichoripetaly | cylindrical | 6.5 | (26) via InfoFlora (https://www.infoflora.ch/) |
| *Symphytum officinale* | tube | high | radial | sympetaly | cylindrical | 15 | (26) via InfoFlora (https://www.infoflora.ch/) |
| *Syringa vulgaris* | disk-tube | high | bilateral | semichoripetaly | cylindrical | 12.5 | FloraWeb (www.floraweb.de) |
| *Taraxacum officinale* | head | low | radial | choripetaly | flat or spherical | 0 | (30) |
| *Taraxacum palustre* | head | low | radial | choripetaly | flat or spherical | NA | |
| *Thymus sp* | gullet | medium | bilateral | semichoripetaly | flat or spherical | 4.5 | (26) via InfoFlora (https://www.infoflora.ch/) |
| *Trifolium pratense* | flag | medium | bilateral | semichoripetaly | cylindrical | 9.67 | (17) via TRY (18) |
| *Trifolium repens* | flag | medium | bilateral | semichoripetaly | flat or spherical | 4.94 | (17) via TRY (18) |
| *Urtica dioica* | disk | low | radial | choripetaly | cylindrical | NA | |
| *Valeriana officinalis* | funnel | low | radial | semichoripetaly | flat or spherical | NA | |
| *Verbena officinalis* | funnel | medium | bilateral | semichoripetaly | cylindrical | 4 | (26) via InfoFlora (https://www.infoflora.ch/) |
| *Veronica chamaedrys* | disk-tube | low | bilateral | semichoripetaly | cylindrical | 1.57 | (17) via TRY (18) |
| *Veronica sp* | disk-tube | low | bilateral | semichoripetaly | cylindrical | NA | |
| *Viburnum opulus* | disk | low | radial | choripetaly | flat or spherical | NA | |
| *Vicia cracca* | flag | high | bilateral | semichoripetaly | cylindrical | 15 | (26) via InfoFlora (https://www.infoflora.ch/) |
| *Vicia lutea* | flag | high | bilateral | semichoripetaly | single | 22.5 | (26) via InfoFlora (https://www.infoflora.ch/) |
| *Vicia sepium* | flag | high | bilateral | semichoripetaly | cylindrical | 16 | (26) via InfoFlora (https://www.infoflora.ch/) |

**Table S6.** Reflectance references. An ID different from NA indicates when data was obtained from the FrED database.

| Plant | FReD ID | Reflectance source | Remarks |
|---|---|---|---|
| *Achillea millefolium* | 1387 | | |
| *Aconitum lycoctonum* | NA | NA | |
| *Aegopodium podagraria* | 1128 | (32) | |
| *Ajuga reptans* | 2110 | (32) | |
| *Alchemilla xanthochlora* | NA | NA | |
| *Alliaria petiolata* | 1578 | (32) | |
| *Angelica sylvestris* | 1808 | | |
| *Anthriscus sylvestris* | NA | NA | |
| *Anthyllis vulneraria* | 2284 | (33) | |
| *Aquilegia atrata* | NA | NA | |
| *Arctium minus* | NA | NA | |
| *Aruncus dioicus* | NA | NA | |
| *Astrantia major* | 3597 | (33) | |
| *Bellis perennis* | 3126 | (33) | |
| *Campanula* sp. | NA | NA | Used the values from *Campanula trachelium* |
| *Campanula trachelium* | 1276 | (32) | |
| *Cardamine impatiens* | NA | NA | |
| *Carduus personata* | NA | NA | |
| *Centaurea jacea* | 4048 | NA | |
| *Centaurea montana* | 3708 | (33) | |
| *Centaurea* sp. | NA | NA | |
| *Chaerophyllum hirsutum* | NA | NA | |
| *Circaea lutetiana* | NA | NA | |
| *Cirsium arvense* | 1416 | (32) | |
| *Cirsium eriophorum* | NA | NA | |
| *Cirsium oleraceum* | 1288 | (33) | |
| *Cirsium palustre* | 1289 | (32) | |
| *Cirsium vulgare* | NA | NA | |
| *Cornus sanguinea* | 1158 | (32) | |
| *Crepis biennis* | NA | NA | |
| *Crepis paludosa* | 1988 | (32) | |
| *Crepis* sp. | NA | NA | Used the values from *Crepis paludosa* |
| *Daucus carota* | 4075 | NA | |
| *Epilobium angustifolium* | NA | (34, 35) | |
| *Epilobium hirsutum* | 1395 | (32) | |
| *Epilobium parviflorum* | 1294 | (32) | |
| *Epilobium* sp. | NA | NA | |
| *Erigeron annuus* | NA | NA | |
| *Eupatorium cannabinum* | 1418 | (32) | |
| *Filipendula ulmaria* | 2130 | (32) | |
| *Fragaria vesca* | 1586 | (32) | |
| *Galeopsis tetrahit* | NA | NA | |

| | | | |
|---|---|---|---|
| *Galium mollugo* | 1160 | (32) | |
| *Galium odoratum* | NA | NA | |
| *Galium sylvaticum* | NA | NA | |
| *Galium verum* | 1396 | (32) | |
| *Geranium pyrenaicum* | 1199 | (33) | |
| *Geranium robertianum* | 1587 | (32) | |
| *Geranium sylvaticum* | NA | NA | |
| *Geum rivale* | 1624 | (33) | |
| *Geum urbanum* | 1978 | (32) | |
| *Glechoma hederacea* | 1663 | (32) | |
| *Heracleum sphondylium* | NA | NA | |
| *Hieracium* sp. | NA | NA | |
| *Hippocrepis comosa* | NA | NA | |
| *Hypericum maculatum* | 1842 | (33) | |
| *Hypericum perforatum* | 1397 | (32) | |
| *Hypochaeris radicata* | NA | (34, 35) | |
| *Impatiens noli-tangere* | 3019 | (32) | |
| *Impatiens parviflora* | 1503 | (32) | |
| *Jacobaea vulgaris* | 1412 | NA | |
| *Knautia dipsacifolia* | 3324 | (33) | |
| *Lamium album* | 1490 | (32) | |
| *Lamium galeobdolon* | 2395 | (33) | |
| *Lamium maculatum* | 1739 | (33) | |
| *Lamium purpureum* | NA | (34, 35) | |
| *Lapsana communis* | 2143 | (32) | |
| *Lathyrus pratensis* | 3508 | (36) | |
| *Leontodon* sp. | NA | NA | Own measurements, used the values from *Leontodon tuberosus* |
| *Leucanthemum vulgare* | 3208 | (33) | |
| *Lonicera ligustrina* | NA | NA | |
| *Lotus corniculatus* | 1424 | (32) | |
| *Malva moschata* | 4234 | NA | |
| *Matricaria discoidea* | 1316 | (33) | |
| *Medicago lupulina* | 2018 | (33) | |
| *Melilotus albus* | NA | NA | |
| *Mentha longifolia* | 1913 | (33) | |
| *Myosotis scorpioides* | NA | NA | |
| *Myosotis* sp. | NA | NA | |
| *Oenothera biennis* | NA | (34, 35) | |
| *Oxalis acetosella* | NA | NA | |
| *Persicaria bistorta* | 3193 | (32) | |
| *Persicaria maculosa* | NA | NA | |
| *Phyteuma spicatum* | NA | NA | |
| *Plantago lanceolata* | NA | NA | |
| *Potentilla erecta* | 1649 | (36) | |

| | | | |
|---|---|---|---|
| *Potentilla reptans* | 1600 | (32) | |
| *Prunella vulgaris* | 1407 | (32) | |
| *Ranunculus* sp. | 1408 | (32) | |
| *Rorippa sylvestris* | NA | NA | |
| *Rubus caesius* | 1409 | (32) | |
| *Rubus fruticosus* | NA | NA | |
| *Rubus idaeus* | NA | NA | |
| *Rubus* sp. | NA | NA | |
| *Salvia glutinosa* | NA | NA | |
| *Salvia pratensis* | 1602 | (32) | |
| *Scorzoneroides autumnalis* | NA | NA | |
| *Silene dioica* | 1251 | (36) | |
| *Silene vulgaris* | 1252 | (36) | |
| *Solidago canadensis* | 3187 | (32) | |
| *Spiraea japonica* | NA | NA | |
| *Stachys alpina* | NA | NA | |
| *Stachys sylvatica* | 1777 | (33) | |
| *Stellaria media* | NA | NA | |
| *Symphoricarpos albus* | 1009 | (33) | |
| *Symphytum officinale* | 2132 | (32) | |
| *Syringa vulgaris* | 1777 | (33) | |
| *Taraxacum officinale* | 1255 | (36) | |
| *Taraxacum palustre* | 1255 | (36) | |
| *Thymus* sp. | 3281 | (33) | Used the same values from *Thymus serpyllum* |
| *Trifolium pratense* | 1654 | (36) | |
| *Trifolium repens* | 1888 | (33) | |
| *Urtica dioica* | NA | NA | |
| *Valeriana officinalis* | 1788 | (33) | |
| *Verbena officinalis* | NA | NA | |
| *Veronica chamaedrys* | 3513 | (32) | |
| *Veronica* sp. | NA | NA | Used the same values from *Veronica chamaedrys* |
| *Viburnum opulus* | 1994 | (32) | |
| *Vicia cracca* | 1257 | (36) | |
| *Vicia lutea* | NA | NA | |
| *Vicia sepium* | 1611 | (32) | |

**Table S7.** Comparison of the observed mean participation coefficient ($\bar{P}$) against the temporal null models. LCL: 95% lower confidence limit, UCL: 95% upper confidence limit, P val: P value.

| Network | Year | Site | $\bar{P}$ | Null model 1 | | | Null model 2 | | | Null model 3 | | |
|---|---|---|---|---|---|---|---|---|---|---|---|---|
| | | | | LCL | UCL | P val | LCL | UCL | P val | LCL | UCL | P val |
| 1 | 2014 | A | 0.302 | 0.484 | 0.591 | < 0.001 | 0.460 | 0.574 | < 0.001 | 0.468 | 0.583 | < 0.001 |
| 2 | 2014 | B | 0.402 | 0.590 | 0.706 | < 0.001 | 0.544 | 0.649 | < 0.001 | 0.538 | 0.643 | < 0.001 |
| 3 | 2014 | C | 0.359 | 0.626 | 0.738 | < 0.001 | 0.557 | 0.653 | < 0.001 | 0.535 | 0.657 | < 0.001 |
| 4 | 2014 | D | 0.345 | 0.522 | 0.625 | < 0.001 | 0.410 | 0.515 | < 0.001 | 0.390 | 0.512 | < 0.001 |
| 5 | 2014 | E | 0.536 | 0.621 | 0.797 | < 0.001 | 0.556 | 0.734 | 0.020 | 0.529 | 0.714 | 0.100 |
| 6 | 2014 | F | 0.494 | 0.600 | 0.771 | < 0.001 | 0.542 | 0.700 | < 0.001 | 0.513 | 0.668 | 0.010 |
| 7 | 2014 | G | 0.313 | 0.516 | 0.660 | < 0.001 | 0.459 | 0.601 | < 0.001 | 0.378 | 0.541 | < 0.001 |
| 8 | 2014 | H | 0.372 | 0.497 | 0.620 | < 0.001 | 0.370 | 0.489 | 0.090 | 0.316 | 0.440 | 0.900 |
| 9 | 2016 | A | 0.067 | 0.249 | 0.452 | < 0.001 | 0.191 | 0.345 | < 0.001 | 0.174 | 0.352 | < 0.001 |
| 10 | 2016 | I | 0.160 | 0.323 | 0.445 | < 0.001 | 0.298 | 0.415 | < 0.001 | 0.284 | 0.410 | < 0.001 |
| 11 | 2016 | K | 0.393 | 0.534 | 0.791 | < 0.001 | 0.511 | 0.778 | < 0.001 | 0.517 | 0.778 | < 0.001 |
| 12 | 2016 | M | 0.310 | 0.628 | 0.769 | < 0.001 | 0.571 | 0.720 | < 0.001 | 0.513 | 0.661 | < 0.001 |
| 13 | 2016 | R | 0.104 | 0.365 | 0.533 | < 0.001 | 0.113 | 0.283 | 0.030 | 0.142 | 0.293 | 0.020 |
| 14 | 2016 | S | 0.088 | 0.438 | 0.599 | < 0.001 | 0.305 | 0.444 | < 0.001 | 0.320 | 0.450 | < 0.001 |

**Table S8.** Comparison of the observed mean versatility ($\bar{V}$) against the temporal null models. LCL: 95% lower confidence limit, UCL: 95% upper confidence limit, P val: P value.

| Network | Year | Site | $\bar{V}$ | Null model 1 | | | Null model 2 | | | Null model 3 | | |
|---|---|---|---|---|---|---|---|---|---|---|---|---|
| | | | | LCL | UCL | P val | LCL | UCL | P val | LCL | UCL | P val |
| 1 | 2014 | A | 0.135 | 0.069 | 0.081 | < 0.001 | 0.072 | 0.088 | < 0.001 | 0.074 | 0.108 | < 0.001 |
| 2 | 2014 | B | 0.241 | 0.100 | 0.118 | < 0.001 | 0.104 | 0.133 | < 0.001 | 0.112 | 0.153 | < 0.001 |
| 3 | 2014 | C | 0.231 | 0.108 | 0.127 | < 0.001 | 0.121 | 0.146 | < 0.001 | 0.121 | 0.172 | < 0.001 |
| 4 | 2014 | D | 0.153 | 0.086 | 0.110 | < 0.001 | 0.103 | 0.133 | < 0.001 | 0.106 | 0.150 | 0.020 |
| 5 | 2014 | E | 0.260 | 0.123 | 0.151 | < 0.001 | 0.138 | 0.169 | < 0.001 | 0.152 | 0.235 | < 0.001 |
| 6 | 2014 | F | 0.327 | 0.120 | 0.157 | < 0.001 | 0.127 | 0.175 | < 0.001 | 0.140 | 0.207 | < 0.001 |
| 7 | 2014 | G | 0.176 | 0.133 | 0.169 | 0.010 | 0.140 | 0.181 | 0.200 | 0.181 | 0.337 | 0.080 |
| 8 | 2014 | H | 0.145 | 0.080 | 0.096 | < 0.001 | 0.090 | 0.111 | < 0.001 | 0.096 | 0.133 | < 0.001 |
| 9 | 2016 | A | 0.131 | 0.111 | 0.132 | 0.040 | 0.114 | 0.160 | 0.950 | 0.131 | 0.207 | 0.080 |
| 10 | 2016 | I | 0.113 | 0.075 | 0.090 | < 0.001 | 0.078 | 0.097 | < 0.001 | 0.101 | 0.157 | 0.440 |
| 11 | 2016 | K | 0.274 | 0.135 | 0.162 | < 0.001 | 0.136 | 0.169 | < 0.001 | 0.125 | 0.169 | < 0.001 |
| 12 | 2016 | M | 0.148 | 0.077 | 0.085 | < 0.001 | 0.084 | 0.095 | < 0.001 | 0.105 | 0.143 | < 0.001 |
| 13 | 2016 | R | 0.217 | 0.104 | 0.128 | < 0.001 | 0.123 | 0.171 | < 0.001 | 0.126 | 0.172 | < 0.001 |
| 14 | 2016 | S | 0.324 | 0.257 | 0.318 | < 0.001 | 0.245 | 0.362 | 0.410 | 0.243 | 0.439 | 0.790 |

**Table S9**. **Robustness model fixed effects and sensitivity analysis.** Fixed-effect estimates from the generalised linear mixed model of $\overline{R50}$ (beta, logit link) fitted to the full dataset and after removing networks sampled in only two of the three diel periods. β, SE, Z, and P(Z) are the coefficient estimate, standard error, Z-statistic, and associated P-value from the model summary. NA values indicate terms that could not be estimated in the reduced subset due to rank deficiency. Values of P < 0.05 are highlighted in bold.

| **Full dataset (42 obs, 14 networks)** | | | | |
|---|---|---|---|---|
| **Predictor** | **β** | **SE** | **Z** | **P (Z)** |
| Mean participation (residuals) | 0.020 | 0.015 | 1.387 | 0.165 |
| Mean versatility | 0.057 | 0.010 | 5.772 | **< 0.001** |
| Mean Participation (Residuals) × Mean versatility | -0.058 | 0.010 | -5.636 | **< 0.001** |
| Plant richness | -0.023 | 0.008 | -2.845 | **0.004** |
| Interaction frequency per network | -0.012 | 0.009 | -1.315 | 0.189 |
| Sampling coverage | 0.047 | 0.009 | 5.086 | **< 0.001** |
| Network layers | 0.080 | 0.008 | 9.804 | **< 0.001** |
| Extinction threshold | 0.011 | < 0.001 | 25.910 | **< 0.001** |
| **Removing afternoon-night-only networks (33 obs, 11 networks)** | | | | |
| Mean participation (residuals) | 0.026 | 0.020 | 1.303 | 0.193 |
| Mean versatility | 0.048 | 0.017 | 2.832 | **0.005** |
| Mean Participation (Residuals) × Mean versatility | -0.053 | 0.012 | -4.476 | **< 0.001** |
| Plant richness | -0.036 | 0.013 | -2.793 | **0.005** |
| Interaction frequency per network | 0.000 | 0.013 | -0.008 | 0.994 |
| Sampling coverage | 0.042 | 0.010 | 4.174 | **< 0.001** |
| Network layers | NA | NA | NA | NA |
| Extinction threshold | 0.011 | < 0.001 | 21.736 | **< 0.001** |

**Table S10**. **Participation model fixed effects and sensitivity analysis.** Fixed-effect estimates from the generalised linear mixed model of plant participation (ordered beta, logit link) fitted to the full dataset and after removing networks sampled in only two of the three diel periods. $\chi^2$, df, and P($\chi^2$) correspond to Type III Wald chi-squared tests, testing the overall significance of each variable. For variables modelled as natural splines (sampling coverage), the $\chi^2$ test evaluates the joint significance of all basis functions. β, SE, Z, and P(Z) are the coefficient estimate, standard error, Z-statistic, and associated P-value from the model summary. NA values indicate terms that could not be estimated in the reduced subset due to rank deficiency. Values of P < 0.05 are highlighted in bold.

| **Full dataset (247 obs, 14 networks)** | | | | | | | |
|---|---|---|---|---|---|---|---|
| Predictor | $\chi^2$ | df | P ($\chi^2$) | β | SE | Z | P(Z) |
| Achromatic contrast | 3.124 | 1 | 0.077 | -0.175 | 0.099 | -1.767 | 0.077 |
| Colour contrast | 1.884 | 1 | 0.17 | -0.14 | 0.102 | -1.372 | 0.170 |
| Floral complexity | 0.525 | 1 | 0.469 | -0.082 | 0.113 | -0.725 | 0.469 |
| Plant height | 7.57 | 1 | **0.006** | 0.288 | 0.105 | 2.751 | **0.006** |
| Floral abundance | 0.683 | 1 | 0.408 | -0.090 | 0.108 | -0.827 | 0.408 |
| Interaction frequency | 9.998 | 2 | **0.007** | | | | |
| spline 1 | | | | -2.091 | 0.737 | -2.838 | **0.005** |
| spline 2 | | | | 0.513 | 0.438 | 1.172 | 0.241 |
| Sampling coverage | 9.785 | 2 | **0.008** | | | | |
| spline 1 | | | | -1.285 | 0.475 | -2.702 | **0.007** |
| spline 2 | | | | -1.540 | 0.528 | -2.914 | **0.004** |
| Network layers | 5.401 | 1 | 0.020 | 0.280 | 0.120 | 2.324 | **0.020** |
| Year | 1 | 0.113 | 0.229 | 0.145 | 1.587 | 0.113 | 0.159 |
| **Remove afternoon-night-only networks (196 obs, 11 networks)** | | | | | | | |
| Predictor | $\chi^2$ | df | P ($\chi^2$) | β | SE | Z | P(Z) |
| Achromatic contrast | 5.214 | 1 | **0.022** | -0.190 | 0.083 | -2.283 | **0.022** |
| Colour contrast | 3.526 | 1 | 0.060 | -0.159 | 0.085 | -1.878 | 0.060 |
| Floral complexity | 0.188 | 1 | 0.665 | -0.04 | 0.093 | -0.433 | 0.665 |
| Plant height | 6.427 | 1 | 0.011 | 0.227 | 0.089 | 2.535 | **0.011** |
| Floral abundance | 2.097 | 1 | 0.148 | -0.144 | 0.100 | -1.448 | 0.148 |
| Interaction frequency | 15.034 | 2 | **0.001** | | | | |
| spline 1 | | | | -1.83 | 0.644 | -2.843 | **0.004** |
| spline 2 | | | | 1.046 | 0.408 | 2.565 | **0.010** |
| Sampling coverage | 8.497 | 2 | **0.014** | | | | |
| spline 1 | | | | -0.966 | 0.366 | -2.636 | **0.008** |
| spline 2 | | | | -1.33 | 0.604 | -2.202 | **0.028** |
| Network layers | NA | NA | NA | NA | NA | NA | |
| Year | 2.977 | 1 | 0.084 | 0.418 | 0.242 | 1.725 | 0.084 |

**Table S11. Versatility model fixed effects and sensitivity analysis.** Fixed-effect estimates from the generalised linear mixed model of plant versatility (ordered beta, logit link) fitted to the full dataset and after removing networks sampled in only two of the three diel periods. $\chi^2$, df, and P($\chi^2$) correspond to Type III Wald chi-squared tests, testing the overall significance of each variable. For variables modelled as natural splines (i.e., all floral traits, floral abundance, and sampling intensity), the $\chi^2$ test evaluates the joint significance of all basis functions. β, SE, Z, and P(Z) are the coefficient estimate, standard error, Z-statistic, and associated P-value from the model summary. NA values indicate terms that could not be estimated in the reduced subset due to rank deficiency. Values of P < 0.05 are highlighted in bold.

| **Full dataset (247 obs, 14 networks)** | | | | | | | |
|---|---|---|---|---|---|---|---|
| Predictor | $\chi^2$ | df | P ($\chi^2$) | β | SE | Z | P(Z) |
| Achromatic contrast | 2.987 | 1 | 0.084 | -0.086 | 0.050 | -1.728 | 0.084 |
| Colour contrast | 7.261 | 2 | **0.027** | | | | |
| spline 1 | | | | -1.545 | 0.581 | -2.658 | **0.008** |
| spline 2 | | | | -0.136 | 0.164 | -0.829 | 0.407 |
| Floral complexity | 13.480 | 2 | **0.001** | | | | |
| spline 1 | | | | 0.039 | 0.227 | 0.171 | 0.864 |
| spline 2 | | | | -0.622 | 0.175 | -3.562 | **< 0.001** |
| Plant height | 9.847 | 2 | **0.007** | | | | |
| spline 1 | | | | -0.502 | 0.434 | -1.157 | 0.247 |
| spline 2 | | | | -0.526 | 0.180 | -2.923 | **0.003** |
| Floral abundance | 151.175 | 3 | **< 0.001** | | | | |
| spline 1 | | | | 1.506 | 0.274 | 5.502 | **< 0.001** |
| spline 2 | | | | 3.514 | 0.945 | 3.717 | **< 0.001** |
| spline 3 | | | | 3.754 | 0.561 | 6.690 | **< 0.001** |
| Interaction frequency | 156.135 | 2 | **< 0.001** | | | | |
| spline 1 | | | | 6.138 | 0.603 | 10.175 | **< 0.001** |
| spline 2 | | | | 3.752 | 1.547 | 2.426 | **0.015** |
| Sampling coverage | 13.281 | 2 | **0.001** | | | | |
| spline 1 | | | | -1.096 | 0.547 | -2.003 | **0.045** |
| spline 2 | | | | -1.566 | 0.431 | -3.637 | **< 0.001** |
| Network layers | 8.552 | 1 | **0.003** | -0.294 | 0.101 | -2.924 | **0.003** |
| Year | 0.001 | 1 | 0.980 | 0.003 | 0.130 | 0.025 | 0.980 |
| **Remove afternoon-night-only networks (196 obs, 11 networks)** | | | | | | | |
| Predictor | $\chi^2$ | df | P ($\chi^2$) | β | SE | Z | P(Z) |
| Achromatic contrast | 1.159 | 1 | 0.282 | -0.052 | 0.048 | -1.076 | 0.282 |
| Colour contrast | 4.856 | 2 | 0.088 | | | | |
| spline 1 | | | | -0.926 | 0.584 | -1.586 | 0.113 |
| spline 2 | | | | 0.195 | 0.162 | 1.206 | 0.228 |
| Floral complexity | 3.577 | 2 | 0.167 | | | | |
| spline 1 | | | | 0.120 | 0.234 | 0.511 | 0.610 |
| spline 2 | | | | -0.311 | 0.184 | -1.687 | 0.092 |
| Plant height | 7.673 | 2 | **0.022** | | | | |
| spline 1 | | | | -0.316 | 0.397 | -0.797 | 0.426 |
| spline 2 | | | | -0.506 | 0.192 | -2.637 | **0.008** |
| Floral abundance | 134.88 | 3 | **< 0.001** | | | | |
| spline 1 | | | | 1.108 | 0.282 | 3.93 | **< 0.001** |

| | | | | | | | |
|---|---|---|---|---|---|---|---|
| spline 2 | | | | 3.229 | 0.898 | 3.594 | **< 0.001** |
| spline 3 | | | | 2.91 | 0.308 | 9.459 | **< 0.001** |
| Interaction frequency | 123.336 | 2 | **< 0.001** | | | | |
| spline 1 | | | | 6.745 | 0.969 | 6.961 | **< 0.001** |
| spline 2 | | | | 6.107 | 2.513 | 2.43 | **0.015** |
| Sampling coverage | 6.084 | 2 | **0.048** | | | | |
| spline 1 | | | | -0.714 | 0.521 | -1.372 | 0.170 |
| spline 2 | | | | -1.882 | 0.782 | -2.406 | 0.016 |
| Network layers | NA | NA | NA | NA | NA | NA | NA |
| Year | 1.716 | 1 | 0.190 | 0.359 | 0.274 | 1.31 | 0.190 |

**Table S12**. Centrality model fixed effects and sensitivity analysis. Fixed-effect results from the centrality model (ordered beta, probit link) refitted on the full dataset and three progressively restricted subsets addressing the unequal diel sampling design. Each subset is labelled with its number of observations and networks. For floral traits interacting with diel period, $\chi^2$ and $P(\chi^2)$ correspond to Type III Wald chi-squared tests on the period × trait interaction term, testing whether the trait effect differs across diel periods; indented sub-rows report the estimated marginal trend (Est.) of each trait within each period, computed via emtrends, with its standard error (SE), Z-statistic, and P-value. For period-invariant covariates, $\chi^2$ tests the main effect and estimates are reported as model coefficients (β); for interaction frequency, modelled as a natural spline, individual basis-function coefficients are given in indented sub-rows. Consistent estimates across subsets indicate that a given effect is robust to the unequal sampling design, whereas effects that change in significance or magnitude across subsets should be interpreted with correspondingly greater caution. Values of $P < 0.05$ are highlighted in bold.

| **Full dataset (450 obs, 22 networks)** | | | | | | | |
|---|---|---|---|---|---|---|---|
| Predictor | χ2 | df | P (χ2) | Est. | SE | Z | P (Z) |
| Achromatic contrast | 12.873 | 2 | **0.002** | | | | |
| Morning | | | | -0.016 | 0.068 | -0.237 | 0.812 |
| Afternoon | | | | -0.038 | 0.047 | -0.807 | 0.420 |
| Night | | | | 0.279 | 0.082 | 3.397 | **0.001** |
| Colour contrast | 5.755 | 2 | 0.056 | | | | |
| Morning | | | | -0.021 | 0.057 | -0.374 | 0.709 |
| Afternoon | | | | 0.027 | 0.045 | 0.601 | 0.548 |
| Night | | | | 0.233 | 0.095 | 2.451 | **0.014** |
| Floral complexity | 1.357 | 2 | 0.507 | | | | |
| Morning | | | | 0.101 | 0.071 | 1.428 | 0.153 |
| Afternoon | | | | 0.061 | 0.049 | 1.25 | 0.211 |
| Night | | | | 0.210 | 0.127 | 1.661 | 0.097 |
| Plant height | 8.505 | 2 | **0.014** | | | | |
| Morning | | | | 0.024 | 0.066 | 0.368 | 0.713 |
| Afternoon | | | | 0.033 | 0.046 | 0.716 | 0.474 |
| Night | | | | 0.408 | 0.128 | 3.193 | **0.001** |
| Floral abundance | 173.357 | 2 | **< 0.001** | | | | |
| Morning | | | | -0.080 | 0.057 | -1.399 | 0.162 |
| Afternoon | | | | -0.018 | 0.051 | -0.364 | 0.716 |
| Night | | | | 0.303 | 0.097 | 3.136 | **0.002** |
| Interaction frequency | 173.357 | 2 | **< 0.001** | | | | |
| spline 1 | | | | 1.897 | 0.171 | 11.119 | **< 0.001** |
| spline 2 | | | | 2.685 | 0.253 | 10.594 | **< 0.001** |
| Sampling coverage | 2.236 | 1 | 0.135 | -0.099 | 0.066 | -1.495 | 0.135 |
| Year | 0.171 | 1 | 0.679 | 0.046 | 0.112 | 0.414 | 0.679 |
| **Remove night-only networks (409 obs, 14 networks)** | | | | | | | |
| Achromatic contrast | 29.092 | 2 | **< 0.001** | | | | |
| Morning | | | | 0.003 | 0.065 | 0.042 | 0.967 |
| Afternoon | | | | -0.025 | 0.044 | -0.568 | 0.570 |
| Night | | | | 0.553 | 0.103 | 5.364 | **< 0.001** |
| Colour contrast | 7.495 | 2 | **0.024** | | | | |

| | | | | | | | |
|---|---|---|---|---|---|---|---|
| Morning | | | | -0.012 | 0.054 | -0.227 | 0.821 |
| Afternoon | | | | 0.031 | 0.042 | 0.746 | 0.456 |
| Night | | | | 0.328 | 0.116 | 2.823 | **0.005** |
| Floral complexity | 2.027 | 2 | 0.363 | | | | |
| Morning | | | | 0.111 | 0.067 | 1.651 | 0.099 |
| Afternoon | | | | 0.066 | 0.046 | 1.422 | 0.155 |
| Night | | | | 0.291 | 0.161 | 1.807 | 0.071 |
| Plant height | 9.489 | 2 | **0.009** | | | | |
| Morning | | | | 0.02 | 0.061 | 0.327 | 0.744 |
| Afternoon | | | | 0.028 | 0.044 | 0.646 | 0.518 |
| Night | | | | 0.469 | 0.142 | 3.29 | **0.001** |
| Floral abundance | 19.482 | 2 | < **0.001** | | | | |
| Morning | | | | -0.072 | 0.055 | -1.31 | 0.19 |
| Afternoon | | | | -0.016 | 0.049 | -0.335 | 0.737 |
| Night | | | | 0.552 | 0.134 | 4.12 | < **0.001** |
| Interaction frequency | 181.75 | 2 | < **0.001** | | | | |
| spline 1 | | | | 1.919 | 0.165 | 11.633 | < **0.001** |
| spline 2 | | | | 2.746 | 0.262 | 10.461 | < **0.001** |
| Sampling coverage | 3.232 | 1 | 0.072 | -0.117 | 0.065 | -1.798 | 0.072 |
| Year | 2.318 | 1 | 0.128 | 0.122 | 0.08 | 1.523 | 0.128 |
| **Remove night-only & morning data (293 obs, 14 networks)** | | | | | | | |
| Achromatic contrast | 28.774 | 1 | < **0.001** | | | | |
| Afternoon | | | | -0.032 | 0.048 | -0.672 | 0.501 |
| Night | | | | 0.569 | 0.112 | 5.064 | < **0.001** |
| Colour contrast | 2.445 | 1 | 0.118 | | | | |
| Afternoon | | | | 0.032 | 0.046 | 0.700 | 0.484 |
| Night | | | | 0.214 | 0.115 | 1.856 | 0.063 |
| Floral complexity | 1.781 | 1 | 0.182 | | | | |
| Afternoon | | | | 0.067 | 0.05 | 1.324 | 0.186 |
| Night | | | | 0.288 | 0.166 | 1.738 | 0.082 |
| Plant height | 6.449 | 1 | **0.011** | | | | |
| Afternoon | | | | 0.030 | 0.048 | 0.628 | 0.530 |
| Night | | | | 0.391 | 0.143 | 2.743 | **0.006** |
| Floral abundance | 18.051 | 1 | < **0.001** | | | | |
| Afternoon | | | | -0.018 | 0.052 | -0.344 | 0.731 |
| Night | | | | 0.535 | 0.128 | 4.165 | < **0.001** |
| Interaction frequency | 123.682 | 2 | < **0.001** | | | | |
| spline 1 | | | | 1.944 | 0.207 | 9.396 | < **0.001** |
| spline 2 | | | | 2.813 | 0.313 | 8.979 | < **0.001** |
| Sampling coverage | 6.221 | 1 | **0.013** | 0.329 | 0.132 | 2.494 | **0.013** |
| Year | 1.966 | 1 | 0.161 | -0.128 | 0.091 | -1.402 | 0.161 |
| **Remove night-only & afternoon-night-only (349 obs, 11 networks)** | | | | | | | |
| Achromatic contrast | 6.35 | 2 | 0.042 | | | | |

| | | | | | | | |
|---|---|---|---|---|---|---|---|
| Morning | | | | -0.015 | 0.062 | -0.239 | 0.811 |
| Afternoon | | | | -0.068 | 0.048 | -1.43 | 0.153 |
| Night | | | | 0.281 | 0.133 | 2.113 | **0.035** |
| Colour contrast | 1.785 | 2 | 0.410 | | | | |
| Morning | | | | -0.002 | 0.053 | -0.033 | 0.974 |
| Afternoon | | | | 0.004 | 0.046 | 0.092 | 0.927 |
| Night | | | | 0.165 | 0.12 | 1.378 | 0.168 |
| Floral complexity | 2.853 | 2 | 0.240 | | | | |
| Morning | | | | 0.093 | 0.065 | 1.431 | 0.152 |
| Afternoon | | | | 0.014 | 0.053 | 0.264 | 0.792 |
| Night | | | | 0.271 | 0.164 | 1.650 | 0.099 |
| Plant height | 0.898 | 2 | 0.638 | | | | |
| Morning | | | | 0.012 | 0.06 | 0.200 | 0.842 |
| Afternoon | | | | 0.021 | 0.046 | 0.450 | 0.653 |
| Night | | | | 0.158 | 0.146 | 1.077 | 0.281 |
| Floral abundance | 4.496 | 2 | 0.106 | | | | |
| Morning | | | | -0.074 | 0.051 | -1.441 | 0.15 |
| Afternoon | | | | -0.026 | 0.049 | -0.526 | 0.599 |
| Night | | | | 0.233 | 0.138 | 1.689 | 0.091 |
| Interaction frequency | 154.386 | 2 | 0 | | | | |
| spline 1 | | | | 1.896 | 0.173 | 10.972 | < **0.001** |
| spline 2 | | | | 2.745 | 0.284 | 9.669 | < **0.001** |
| Sampling coverage | 3.392 | 1 | 0.066 | -0.119 | 0.065 | -1.842 | 0.066 |
| Year | 0.05 | 1 | 0.822 | -0.023 | 0.104 | -0.225 | 0.822 |

**Table S13.** Pairwise contrasts of period-specific trait effects on centrality. Contrasts compare the estimated marginal trend (slope) of each floral trait between diel periods, derived from the centrality model via emtrends followed by pairwise comparison. Each row gives the difference (Est. diff) in the trait's slope between two periods (e.g., Afternoon − Night), with its standard error (SE), Z-statistic, and P-value; P-values are adjusted for multiple comparisons using the Tukey method. A significant contrast indicates that the strength of a trait's association with centrality differs significantly between the two diel periods, providing a formal test of the diel dependence of that trait's effect.

| **Full dataset (448 obs, 22 networks)** | | | | | |
|---|---|---|---|---|---|
| Predictor | Contrast | Est. diff | SE | Z | P |
| Achromatic contrast | Afternoon − Morning | -0.010 | 0.074 | -0.132 | 0.990 |
| | Afternoon − Night | -0.318 | 0.089 | -3.568 | **0.001** |
| | Morning − Night | -0.309 | 0.103 | -3.004 | **0.008** |
| Colour contrast | Afternoon − Morning | 0.075 | 0.068 | 1.106 | 0.510 |
| | Afternoon − Night | -0.205 | 0.099 | -2.069 | 0.096 |
| | Morning − Night | -0.28 | 0.107 | -2.615 | **0.024** |
| Floral complexity | Afternoon − Morning | -0.011 | 0.081 | -0.137 | 0.990 |
| | Afternoon − Night | -0.149 | 0.133 | -1.124 | 0.499 |
| | Morning − Night | -0.138 | 0.143 | -0.967 | 0.598 |
| Plant height | Afternoon − Morning | 0.035 | 0.074 | 0.476 | 0.882 |
| | Afternoon − Night | -0.370 | 0.130 | -2.837 | **0.013** |
| | Morning − Night | -0.406 | 0.140 | -2.906 | **0.010** |
| Floral abundance | Afternoon − Morning | 0.072 | 0.069 | 1.044 | 0.549 |
| | Afternoon − Night | -0.319 | 0.102 | -3.108 | **0.005** |
| | Morning − Night | -0.391 | 0.108 | -3.600 | **0.001** |
| **Remove night-only networks (407 obs, 14 networks)** | | | | | |
| Achromatic contrast | Afternoon − Morning | -0.016 | 0.073 | -0.223 | 0.973 |
| | Afternoon − Night | -0.579 | 0.108 | -5.347 | **< 0.001** |
| | Morning − Night | -0.563 | 0.119 | -4.753 | **< 0.001** |
| Colour contrast | Afternoon − Morning | 0.069 | 0.066 | 1.04 | 0.552 |
| | Afternoon − Night | -0.292 | 0.119 | -2.459 | **0.037** |
| | Morning − Night | -0.361 | 0.125 | -2.900 | **0.010** |
| Floral complexity | Afternoon − Morning | -0.017 | 0.079 | -0.209 | 0.976 |
| | Afternoon − Night | -0.227 | 0.165 | -1.373 | 0.355 |
| | Morning − Night | -0.210 | 0.173 | -1.212 | 0.446 |
| Plant height | Afternoon − Morning | 0.034 | 0.072 | 0.478 | 0.882 |
| | Afternoon − Night | -0.436 | 0.144 | -3.016 | **0.007** |
| | Morning − Night | -0.470 | 0.152 | -3.085 | **0.006** |
| Floral abundance | Afternoon − Morning | 0.065 | 0.067 | 0.973 | 0.594 |
| | Afternoon − Night | -0.566 | 0.138 | -4.100 | **< 0.001** |
| | Morning − Night | -0.632 | 0.142 | -4.440 | **< 0.001** |
| **Remove night-only & morning data (293 obs, 14 networks)** | | | | | |
| Achromatic contrast | Afternoon − Night | -0.602 | 0.111 | -5.419 | **< 0.001** |
| Colour contrast | Afternoon − Night | -0.17 | 0.113 | -1.499 | 0.134 |
| Floral complexity | Afternoon − Night | -0.218 | 0.164 | -1.334 | 0.182 |
| Plant height | Afternoon − Night | -0.356 | 0.143 | -2.487 | **0.013** |

| | | | | | |
|---|---|---|---|---|---|
| Floral abundance | Afternoon − Night | -0.551 | 0.131 | -4.216 | < **0.001** |
| **Remove night-only & afternoon-night-only (347 obs, 11 networks)** | | | | | |
| Achromatic contrast | Afternoon − Morning | -0.043 | 0.075 | -0.579 | 0.831 |
| | Afternoon − Night | -0.354 | 0.139 | -2.547 | **0.029** |
| | Morning − Night | -0.310 | 0.144 | -2.159 | 0.078 |
| Colour contrast | Afternoon − Morning | 0.035 | 0.067 | 0.514 | 0.864 |
| | Afternoon − Night | -0.162 | 0.123 | -1.311 | 0.389 |
| | Morning − Night | -0.196 | 0.129 | -1.524 | 0.280 |
| Floral complexity | Afternoon − Morning | -0.052 | 0.082 | -0.631 | 0.803 |
| | Afternoon − Night | -0.261 | 0.17 | -1.541 | 0.272 |
| | Morning − Night | -0.210 | 0.176 | -1.193 | 0.457 |
| Plant height | Afternoon − Morning | 0.035 | 0.073 | 0.471 | 0.885 |
| | Afternoon − Night | -0.138 | 0.151 | -0.918 | 0.629 |
| | Morning − Night | -0.173 | 0.156 | -1.109 | 0.508 |
| Floral abundance | Afternoon − Morning | 0.057 | 0.066 | 0.858 | 0.667 |
| | Afternoon − Night | -0.258 | 0.144 | -1.797 | 0.170 |
| | Morning − Night | -0.315 | 0.145 | -2.169 | 0.077 |

### References

1. C. Caron, P. Homewood, W. Wildi, The original Swiss flysch: a reappraisal of the type deposits in the Swiss prealps. *Earth-Science Reviews* **26**, 1–45 (1989).

2. S. Stefani, *et al.*, Genetic divergence of Clematis alpina in the Swiss Prealps: a tale of the margins. *Alp Botany* **135**, 167–185 (2025).

3. R. H. Gibson, B. Knott, T. Eberlein, J. Memmott, Sampling method influences the structure of plant–pollinator networks. *Oikos* **120**, 822–831 (2011).

4. A. Stefanaki, A. Kantsa, T. Tscheulin, M. Charitonidou, T. Petanidou, Lessons from Red Data Books: Plant Vulnerability Increases with Floral Complexity. *PLOS ONE* **10**, e0138414 (2015).

5. N. Frydman, S. Freilikhman, I. Talpaz, S. Pilosof, Practical guidelines and the EMLN R package for handling ecological multilayer networks. *Methods Ecol. Evol.* **14**, 2964–2973 (2023).

6. M. C. Hutchinson, *et al.*, Seeing the forest for the trees: Putting multilayer networks to work for community ecology. *Funct. Ecol.* **33**, 206–217 (2019).

7. B. Padrón, M. Nogales, A. Traveset, Alternative approaches of transforming bimodal into unimodal mutualistic networks. The usefulness of preserving weighted information. *Basic Appl. Ecol.* **12**, 713–721 (2011).

8. S. Hervías-Parejo, *et al.*, Spatio-temporal variation in plant–pollinator interactions: a multilayer network approach. *Oikos* **2023**, e09818 (2023).

9. T. Opsahl, "Structure and Evolution of Weighted Networks," University of London (Queen Mary College), London, UK. (2009).

10. M. E. J. Newman, The structure of scientific collaboration networks. *Proc. Natl. Acad. Sci. U.S.A.* **98**, 404–409 (2001).

11. H. W. Krenn, J. D. Plant, N. U. Szucsich, Mouthparts of flower-visiting insects. *Arthropod Struct. Dev.* **34**, 1–40 (2005).

12. H. W. Krenn, "Form and Function of Insect Mouthparts" in *Insect Mouthparts: Form, Function, Development and Performance*, H. W. Krenn, Ed. (Springer International Publishing, 2019), pp. 9–46.

13. F. Karolyi, "What's on the Menu: Floral Tissue, Pollen or Nectar? Mouthpart Adaptations of Anthophilous Beetles to Floral Food Sources" in *Insect Mouthparts: Form, Function, Development and Performance*, H. W. Krenn, Ed. (Springer International Publishing, 2019), pp. 419–442.

14. C. W. Schaefer, A. R. Panizzi, Eds., *Heteroptera of Economic Importance* (CRC Press, 2000).

15. P. G. Kevan, H. G. Baker, Insects as Flower Visitors and Pollinators. *Annual Review of Entomology* **28**, 407–453 (1983).

16. G. Bockwinkel, K.-P. Sauer, Panorpa scorpionflies foraging in spider webs — kleptoparasitism at low risk. *Bull. Br. arachnol. Soc.* **9**, 110–112 (1993).

17. R. R. Junker, M. H. Lechleitner, J. Kuppler, L.-M. Ohler, Interconnectedness of the Grinnellian and Eltonian Niche in Regional and Local Plant-Pollinator Communities. *Front. Plant Sci.* **10** (2019).

18. J. Kattge, *et al.*, TRY plant trait database – enhanced coverage and open access. *Glob. Change Biol.* **26**, 119–188 (2020).

19. S. Anton, B. Denisow, K. Milaniuk, Flowering, pollen production and insect visitation in two Aconitum species (Ranunculaceae). *Acta Agrobot.* **67** (2014).

20. K. Winkler, F. L. Wäckers, L. V. Kaufman, V. Larraz, J. C. Van Lenteren, Nectar exploitation by herbivores and their parasitoids is a function of flower species and relative humidity. *Biol. control* **50**, 299–306 (2009).

21. P. B. Cavers, M. I. Heagy, R. F. Kokron, The biology of Canadian weeds.: 35. *Alliaria petiolata* (M. Bieb.) Cavara and Grande. *Can. J. Plant Sci.* **59**, 217–229 (1979).

22. F. S. Gilbert, "Morphological and foraging ecology of hoverflies (Diptera: Syrphidae)," University of Cambridge, Cambridge. (1981).

23. F. Baden-Böhm, M. App, J. Thiele, The FloRes Database: A floral resources trait database for pollinator habitat-assessment generated by a multistep workflow. *Biodivers. Data J.* **10**, e83523 (2022).

24. E. Puidet, J. Liira, J. Paal, M. Pärtel, S. Pihu, Morphological variation in eight taxa of Anthyllis vulneraria s. lato (Fabaceae). *Ann. Bot. Fenn.* **42**, 293–304 (2005).

25. M. A. Becher, *et al.*, Bumble-BEEHAVE: A systems model for exploring multifactorial causes of bumblebee decline at individual, colony, population and community level. *J. Appl. Ecol.* **55**, 2790–2801 (2018).

26. K. Lauber, G. Wagner, A. Gygax, *Flora Helvetica – Illustrated Flora of Switzerland*, 7th Ed. (Haupt Verlag, 2018).

27. B. Denisow, M. Strzałkowska-Abramek, M. Bożek, A. Jeżak, Ornamental Representatives of the Genus Centaurea L. as a Pollen Source for Bee Friendly Gardens. *J. Apic. Sci.* **58**, 49–58 (2014).

28. P. C. J. van Rijn, F. L. Wäckers, Nectar accessibility determines fitness, flower choice and abundance of hoverflies that provide natural pest control. *Journal of Applied Ecology* **53**, 925–933 (2016).

29. W. S. Armbruster, D. A. Guinn, The Solitary Bee Fauna (Hymenoptera: Apoidea) of Interior and Arctic Alaska: Flower Associations, Habitat Use, and Phenology. *J. Kans. Entomol. Soc.* **62**, 468–483 (1989).

30. S. A. Corbet, Butterfly nectaring flowers: butterfly morphology and flower form. *Entomol. Exp. Appl.* **96**, 289–298 (2000).

31. M. Bieniasz, E. Dziedzic, G. Słowik, Biological features of flowers influence the fertility of Lonicera spp. cultivars. *Hortic. Environ. Biotechnol.* **60**, 155–166 (2019).

32. A. Gumbert, J. Kunze, L. Chittka, Floral colour diversity in plant communities, bee colour space and a null model. *Proceedings of the Royal Society of London. Series B: Biological Sciences* **266**, 1711–1716 (1999).

33. L. Chittka, A. Shmida, N. Troje, R. Menzel, Ultraviolet as a component of flower reflections, and the colour perception of hymenoptera. *Vision Research* **34**, 1489–1508 (1994).

34. C. J. van der Kooi, I. Pen, M. Staal, D. G. Stavenga, J. T. M. Elzenga, Competition for pollinators and intra-communal spectral dissimilarity of flowers. *Plant Biol J* **18**, 56–62 (2016).

35. C. J. van der Kooi, J. T. M. Elzenga, M. Staal, D. G. Stavenga, How to colour a flower: on the optical principles of flower coloration. *Proc. R. Soc. B* **283**, 20160429 (2016).

36. L. Chittka, Optimal Sets of Color Receptors and Color Opponent Systems for Coding of Natural Objects in Insect Vision. *Journal of Theoretical Biology* **181**, 179–196 (1996).
